# Regulation of *Cyclin E* by transcription factors of the naïve pluripotency network in mouse embryonic stem cells

**DOI:** 10.1101/623884

**Authors:** Fabrice Gonnot, Diana Langer, Pierre-Yves Bourillot, Nathalie Doerflinger, Pierre Savatier

## Abstract

Continuous, non-cell cycle-dependent expression of cyclin E is a characteristic feature of mouse embryonic stem cells (ESCs). We studied the 5’ regulatory region of *Cyclin E*, also known as *Ccne1*, and identified binding sites for transcription factors of the naïve pluripotency network, including Esrrb, Klf4, and Tfcp2l1 within 1 kilobase upstream of the transcription start site. Luciferase assay and chromatin immunoprecipitation-quantitative polymerase chain reaction (ChiP–qPCR) study highlighted one binding site for Esrrb that is essential to transcriptional activity of the promoter region, and three binding sites for Klf4 and Tfcp2l1. Knockdown of Esrrb, Klf4, and Tfcp2l1 reduced *Cyclin E* expression whereas overexpression of Esrrb and Klf4 increased it, indicating a strong correlation between the expression level of these factors and that of cyclin E. We observed that cyclin E overexpression delays differentiation induced by Esrrb depletion, suggesting that cyclin E is an important target of Esrrb for differentiation blockade. We observed that mESCs express a low level of miR-15a and that transfection of a miR-15a mimic decreases *Cyclin E* mRNA level. These results lead to the conclusion that the high expression level of *Cyclin E* in mESCs can be attributed to transcriptional activation by Esrrb as well as to the absence of its negative regulator, miR-15a.

## Introduction

Cyclin E is a regulatory subunit of cyclin-dependent kinase (Cdk) 2 involved in many cellular processes including cell cycle progression, replication complex assembly, centrosome cycle, and epigenetic regulation. Its expression is regulated at both the transcriptional and protein level to achieve a timely control of cell division in connection with cell environment and fate decision. Deregulated expression of cyclin E has been shown to play a key role in tumorigenesis [1, 2]. Transcription of the *Cyclin E* gene, also known as *Ccne1*, is activated during the G_1_ phase and depends on mitogenic input, which is integrated through E2F and Myc transcription factors [3, 4]. E2F activity is regulated by phosphorylation of the retinoblastoma (Rb) protein in response to cyclin D/Cdk4 and cyclin D/Cdk6 kinase activity [5, 6]. The miRNA miR-15a was shown to act as a negative regulator of *Cyclin E* in somatic cells [7, 8]. Since, both *Cyclin E* and *mir-15a* are direct transcriptional targets of E2F, it raises the possibility that E2F, miR-15, and cyclin E constitute a feed-forward loop that modulates E2F activity and cell-cycle progression [8].

There is a growing body of evidence showing that the cell cycle of mouse embryonic stem cells (ESCs) lacks some of the regulatory pathways that operate in somatic cells [9–11]. These include extensive phosphorylation of the Rb family proteins despite little cyclin D/Cdk4 kinase activity [12], p16^ink4a^-resistant residual cyclin D3/Cdk6 kinase activity [13], and lack of functional Chk/p53/p21^cip1^ and Chk/Cdc25A pathways resulting in the absence of the DNA damage checkpoint in the G_1_ phase [14–16]. A key feature of the pluripotent stem cell cycle is the constitutive activity of Cdk2 due to seemingly continuous expression of both cyclin E and A throughout the cell cycle [17, 18] in addition to low expression levels of the Cdk2 inhibitors p21^cip1^, p27^kip1^, and p57^kip2^ [12, 17]. In a previous report, we showed that cyclin E partially rescues ESC differentiation induced by leukemia inhibitory factor (LIF) starvation, suggesting that cyclin E participates in the regulation of pluripotency [19]. It was established that cyclin E:Cdk2 complexes phosphorylate and thereby stabilize the core pluripotency factors Nanog, Sox2, and Oct4 [20]. These findings point to a connection between the cell cycle machinery regulating G_1_/S phase transition and the core pluripotency network [21].

In this context, it is important to understand how *Cyclin E* is transcriptionally regulated in pluripotent stem cells. We hypothesized that the transcription factors of the naïve pluripotency network would participate in the transcriptional regulation of *Cyclin E*. These factors include the cardinal pluripotency factors Oct4, Sox2, and Nanog, as well as the ancillary transcription factors Klf2, Klf4, Klf5, Esrrb, Tbx3, Gbx2, Nr0b1, and Tfcp2l1, all of which have been shown to sustain the naive state of pluripotency in mice [22–34]. The present study points to Esrrb as a transcriptional activator and miR-15a as a negative regulator of *Cyclin E* in ESCs.

## Material and Methods

### In silico analysis

Published data were obtained from *NCBI Gene Expression Omnibus* (http://www.ncbi.nlm.nih.gov/geo) and analyzed using *UCSC Genome Browser* ([35]; http://genome.ucsc.edu). DNAse I hypersensitive sites, were identified from GSM1003830 (DNAseDgf on mESC-CJ7), GSM1014154 (DNAseHS on mESC-E14), and GSM1014187 (DNAseHS on mESC-CJ7) datasets. Histone marks were identified from GSM769008 (H3K4me3 on mESC-Bruce4), GSM1000089 (H3K27me3 on mESC-Bruce4) and GSM1000124 (H3K4me3 on mESC-E14) datasets. ChIP-seq data were from GSM288345 (Nanog), GSM288346 (Oct4), GSM288347 (Sox2), GSM288349 (E2f1), GSM288350 (Tfcp2I1), GSM288353 (Stat3), GSM288354 (Klf4), GSM288355 (Esrrb), and GSM288356 (c-Myc) compendiums [36], and GSM470523 (Nr5a2) [37] and GSM1208217 (Klf4) [38]. Several resources were used to predict the transcription factor binding site (TFBS)’s relative scores on the genomic sequence upstream of the *Ccne1* gene, downloaded from the *Ensembl* database (genome assembly GRCm38/mm10, December 2011). They include *JASPAR* [[39]; http://jaspar.genereg.net], *TRANSFAC 7.0 public by BIOBASE* [[40]; http://www.gene-regulation.com], *MAPPER2* [[41]; http://genome.ufl.edu/mapperdb], *cisRED mouse v4* [[42]; http://www.cisred.org/mouse4], *UniPROBE* [[43]; http://the_brain.bwh.harvard.edu/uniprobe], *MotifViz* [[44]; http://biowulf.bu.edu/MotifViz] and *CONSITE* [[45]; http://consite.genereg.net]. A transcription factor and DNA sequence matching degree greater than 80% was considered as a putative TFBS.

### Quantitative real-time PCR (qRT-PCR)

Total RNA was isolated from cell pellets using TRIzol (Ambion) according to the manufacturer’s protocol and reverse-transcribed using a High-Capacity RNA-to-cDNA kit (Applied Biosystems). For microRNAs reverse-transcription, a stem-loop primer specific to each miRNA was used. Real-time PCR was performed using the StepOnePlus real-time PCR system (Applied Biosystems) and Fast SBYR Green Master Mix (Applied Biosystems) according to the manufacturer’s instructions. The relative quantitation of gene expression was calculated using StepOne Software 2.3 (Applied Biosystems). Expression of the target genes was normalized to those of the mouse *β-actin* gene (*Actb*) or to the mouse *sno234* RNA for miRNA. Primers are listed in **Table S1**.

### ChIP-PCR

ChIP for Esrrb, Klf4, and Tfcp2l1 was performed on E14Tg2a ESCs using previously described protocols [46]. In brief, 10^7^ cells were cross-linked with 1% formaldehyde for 15 min. Chromatin was sonicated to a length of less than 400 bp, and subsequently immunoprecipitated with 5 μg of anti-Esrrb (Perseus, pp-H6705-00), anti-Klf4 (Stemgent, 09-0021), and anti-Tfcp2l1 (AbCam, ab123354). DNA fragments encompassing binding sites for Esrrb, Klf4, and Tfcp2l1 in the P region of *Cyclin E* and the *Nanog* promoters were subsequently amplified by PCR. A 3’ untranslated region of the *Cyclin E* gene lacking putative binding sites for Esrrb, Klf4, and Tfcp2l1 was used as negative control. Primers are listed in **Table S2**. ChIP-qPCR data obtained for each specific antibody were normalized using the percent input method that normalizes according to the amount of chromatin input. The percentage value for each sample was calculated based on the equation as follows: % Input =100 × [primer pair efficiency]^(Ct[adjusted input] - Ct[IP]). The “% Input” value represents the enrichment of factor on specific region.

### Plasmid constructs

Regions I (P) (1.5 kb), II (PE) (1.5 kb), and III (DE) (1.7 kb) of the *Cyclin E* gene 5’ flanking sequence were synthesized by GeneArt (Invitrogen) with appropriate restriction sites at both ends and subcloned into the pMA plasmid to generate *pMA-P, pMA-PE*, and *pMA-DE* plasmids, respectively (**Table S3**). A 1,512 base pair (bp) BglII-HindIII fragment encompassing region P was prepared from *pMA-P* and subcloned between BglII and Hind III in *pGL4.10[luc2]* (Promega) to generate *pGL4.10-P*. A 1,506 bp MluI fragment encompassing region PE was prepared from *pMA-PE* and subcloned into the MluI site in *pGL4.10-P* to generate *pGL4.10-P+PE*. A 1,706 bp EcoRV-BglII fragment was subcloned between EcoRV and BglII sites in *pGL4.10-P* and *pGL4.10-P+PE* to generate *pGL4.10-P+DE* and *pGL4.10-P+PE+DE*, respectively.

For site-directed mutagenesis of *Esrrb, Klf4*, and *Tfcp2l1* binding sites, the *pGL4.10-P* plasmid was mutated by PCR using mutant primers and Q5 site-directed mutagenesis kit (New England Biolabs, E0554) according to the manufacturer’s instructions. Mutant primers were designed to modify 10–14 bp encompassing *Esrrb, Klf4*, and *Tfcp2l1* binding sites into 10–14 bp sequences with low binding scores using JASPAR software (**Table S4**).

The shRNA sequences used to knock down *Esrrb, Klf4*, and *Tfcp2l1* are described in **Table S5**. An expression cassette containing the *eGFP* reporter was synthesized by GeneArt (Invitrogen) with MfeI restriction sites at either ends and subcloned into pMA to generate *pMA–eGFP-miR30* plasmid as previously described [47]. shRNA sequences for *Esrrb, Klf4, Tfcp2l1*, and a control sequence were subsequently introduced between XhoI and EcoRI restriction sites in *pMA–eGFP–miR30* to generate *pMA–eGFP–miR30–shEsrrb, pMA–eGFP–miR30–shKlf4, pMA–eGFP–miR30–shTfcp2l1*, and *pMA–eGFP–miR30–shControl*, respectively. MfeI fragments containing the seven *eGFP–miR30–shRNA* sequences were subsequently subcloned into the EcoRI site in *pBS31* [48] to generate *pBS31-TetON–shEsrrb#1, pBS31-TetON–shEsrrb#2, pBS31–TetON–shKlf4#1, pBS31–TetON–shKlf4#2, pBS31–TetON–shTfcp2l1#1, pBS31–TetON–shTfcp2l1#2*, and *pBS31–TetON–shControl*, respectively.

For cDNA overexpression, the coding sequences of mouse *Esrrb (NM_011934), Klf4 (NM_010637)*, and *Tfcp2l1 (NM_023755)* were amplified by PCR with Q5 DNA polymerase (NEB, M0493) and primers containing an EcoRI restriction site and subsequently subcloned into the unique EcoRI restriction site in pBS31 [48] to generate*pBS31–TetON-Esrrb, pBS31– TetONhKlf4*, and *pBS31–TetON–Tfcp2l1*. For *Tfcp2l1*, before pBS31 subcloning, the PCR product was cloned into pJET1.2 with the CloneJET PCR Cloning Kit (Thermo Scientific, K1231), according to the manufacturer’s instructions, and a silent mutation of the EcoRI site, present at position 523-528 of the sequence was performed (GAATTC in to GAGTTC) according to the previously described protocol.

### Cell culture, generation of stable transfectants, and colony assay

Parental E14Tg2a, E14Tg2a–Fucci, KH2 [47], and EKOiE [49] ESC lines were routinely cultured on 0.1% gelatin-coated dishes in Glasgow’s Modified Eagle Medium supplemented with 10% fetal calf serum, 100 μM nonessential amino acids, 100 U/mL penicillin, 100 μg/mL streptomycin, 2 mM L-glutamine, 1 mM sodium pyruvate, 100 βM 2-mercaptoethanol, and 1000 U/mL LIF. Routine culture of EKOiE ESCs included 1 μg/mL Doxycyclinz [49]. Differentiation of E14Tg2a–Fucci cells was induced after withdrawal of LIF for 5 days. Protocol for routine culture of EpiSCsis described elsewhere [50].

For generation of stable transfectants, 1 × 10^6^ KH2 ESCs were electroporated with 5 μg of *pBS31–TetON* plasmid and 5 μg of *pCAGgs–FLPe* plasmid using the Neon system (Invitrogen) with two impulses (20 ms, 1300 volts). After 48 h, stable transfectants were selected with 40 μg/mL Hygromycin B (Roche, 10843555001). cDNA and shRNA expression were induced by Doxycycline (Sigma, D9891) at concentrations ranging from 0.1 to 1.0 μg/mL. EKOiE-^Myc^Cyclin E and EKOiE–control cells were generated by electroporating EKOiE cells with 10 μg of *pCAGgs*–^Myc^Cyclin E plasmid [19] followed by Hygromycin selection.

For colony assays, ESCs were plated at a density of 10^3^ cells per gelatin-coated 100-mm tissue culture dish in complete ESC medium. Cells were exposed to the medium without doxycycline for 1 to 7 days. The protocol for *in situ* detection of alkaline phosphatase activity is described elsewhere [19].

### Infection with lentivirus vectors, flow cytometry, cell transfection and luciferase assay

Production of simian immunodeficiency virus (SIV)-derived lentivectors expressing the Fucci reporters *mKO2:Cdt1, mAG:Geminin* and infection of mouse ESCs are described elsewhere [19]. Cells were either analyzed using a LSRFortessa X-20 (Becton-Dickinson), or sorted with a FACSAria cell sorter (Becton-Dickinson) as described in [19]. For transient expression assay, 5 × 10^4^ E14Tg2a cells were transfected with 100 ng of reporter plasmids (*pGL4.10*[*luc2*] and their derivatives) and 1 ng of *pGL4.70*[*hRluc*] control plasmid using Lipofectamine 2000 (Invitrogen) in 96-well plates. Luciferase activity was measured after 48 h using the *Dual-Glo Luciferase Assay System* kit (Promega) and the *GloMax Multi Detection System* (Promega). For microRNAs mimics transient expression assay, 25 mM of miR-15a-5p (Ambion, 4464066 - MC10235) and miR-1 positive control (Ambion, 4464065) mimics were transfected and the gene expression was measured after 48 h.

### Immunoblotting and immunolabeling

For immunoblotting, frozen cell pellets were lysed in RIPA buffer complemented with protease and phosphatase inhibitors. Protein lysates were then cleared by centrifugation (17,000 × *g* for 20 min). After SDS-PAGE and electroblotting on polyvinylidene fluoride, the membranes were incubated with specific primary antibodies (mouse anti-Esrrb, Perseus PP-H6705-00; rabbit anti-Klf4, Santa Cruz, sc-20691; rabbit anti-Tfcp2l1, AbCam, ab123354; anti-βactin, Sigma, A3854). Blots were incubated with horseradish peroxidase (HRP)-coupled sheep anti-mouse IgG (GE Healthcare, NA931VS) and (HRP)-coupled goat anti-rabbit IgG (GE Healthcare, NA934VS), and developed with Clarity Western ECL Substrate (Bio-Rad, 1705060).

For immunolabeling, cells were fixed with 2% paraformaldehyde in PBS at 4°C for 20 min, and permeabilized in Tris-buffered saline (TBS; 50 mM Tris [pH 7.6], 0.9% NaCl, and 0.2% Triton X-100). The cells were then incubated overnight at 4°C with primary antibodies [anti-cyclin E1 rabbit polyclonal, Santa Cruz, sc-481 (1/100 dilution); anti-mKO2 mouse IgG1 (1/200 dilution), Clinisciences, M168-3M; anti-Esrrb mouse IgG2a (1/500 dilution), Perseus, PP-H6705-00; anti-Klf4 mouse IgG1κ (1/100 dilution), Stemgent, 09-0021; anti-Oct4 rabbit polyclonal (1/300 dilution), Santa Cruz, sc-9081; anti-Oct4 mouse IgG2b (1/300 dilution), Santa Cruz, sc-5279; anti-SOX2 mouse IgG2a (1/50), R&D Systems, MAB2018]. After three rinses (10 min each) with TBS, the cells were incubated with fluorochrome-conjugated secondary antibody [Alexa Fluor 488-conjugated donkey anti-rabbit IgG [H + L], (1/500 dilution), Life Technologies, A21206; Alexa Fluor 647-conjugated donkey anti-mouse IgG [H + L], (1/400 dilution), Life Technologies, A31571] at room temperature for 1 h. The cells were examined under confocal imaging (DM 6000 CS SP5; Leica). Acquisitions were performed using an oil immersion objective (40×/1.25 0.75, PL APO HCX; Leica).

## Results

### Cell cycle expression patterns of cyclins and pluripotency factors

Expression patterns of *Cyclin E* (*Ccne1*), *Cyclin E2* (*Ccne2*), and *Cyclin A* (*Ccna1*) were examined during differentiation of ESCs into embryoid bodies, and compared with that of transcription factors implicated in the regulation of pluripotency including *Oct4, Sox2, Nanog, Esrrb, Klf2, Klf4, Klf5*, and *Tfcp2l1*. Only *Cyclin E* mRNA decreased during differentiation concomitantly with mRNA of all the transcription factors analyzed (**Fig. 1A**). We next examined the expression of pattern for *Cyclin E, Cyclin E2*, and *Cyclin A*, and for 10 pluripotency genes (*Oct4*, *Sox2, Nanog, Esrrb, Klf2, Klf4, Gbx2, Tfcp2l1, Stat3* and *Nr5a2)* in each phase of the cell cycle in ESCs using the Fucci reporter to sort cells out according to their position in the cell cycle (**Fig. 1B**). No significant variation in mRNA levels for *Cyclin E* and for the 10 pluripotency genes was observed between mAG(−)/mKO2(−) (G_1_ phase), mAG(+)/mKO2(−) (S phase), and mAG(++)/mKO2(−) (G_2_ phase) (**Fig. 1C**). In line with expression patterns described in somatic cells, *Cyclin E2* and *Cyclin A* showed lower and higher expression in the G_2_ phase, respectively. Note that the mAG(−)/mKO2(+) fraction (cells in the late G_1_ phase) were excluded from analysis as most of them displayed low or no expression of Oct4 (**Suppl. Fig. 1A**), and therefore have spontaneously committed to differentiation. Co-expression of cyclin E and the pluripotency regulators Oct4, Esrrb and Klf4 was confirmed by immunofluorescence analysis. Rare Oct4-negative, Esrrb-negative, Sox2-negative, and Klf4-negative cells showed low cyclin E content in line with downregulation of *Cyclin E* transcripts observed during controlled differentiation (**Suppl. Fig. 1B**). After differentiation induced by withdrawal of LIF for 5 days, *cyclin E* transcripts displayed a somatic-like pattern of cell cycle expression, showing higher levels in G_1_ and S phase with respect to the G_2_ phase (**Fig. 1D**). Examination of transcripts levels in a whole cell population of EpiSC revealed both a strong reduction for all naïve pluripotency markers and a 70% reduction for *Cyclin E* (**Figure 1E**). Taken together these results indicate that naïve ES cells express *Cyclin E* mRNAs at a constant level throughout the cell-cycle. *Cyclin E* expression is down-regulated in pluripotent stem cells in the primed state of pluripotency. It is further reduced in differentiated cells, where it resumes cyclic expression.

**Figure 1:**
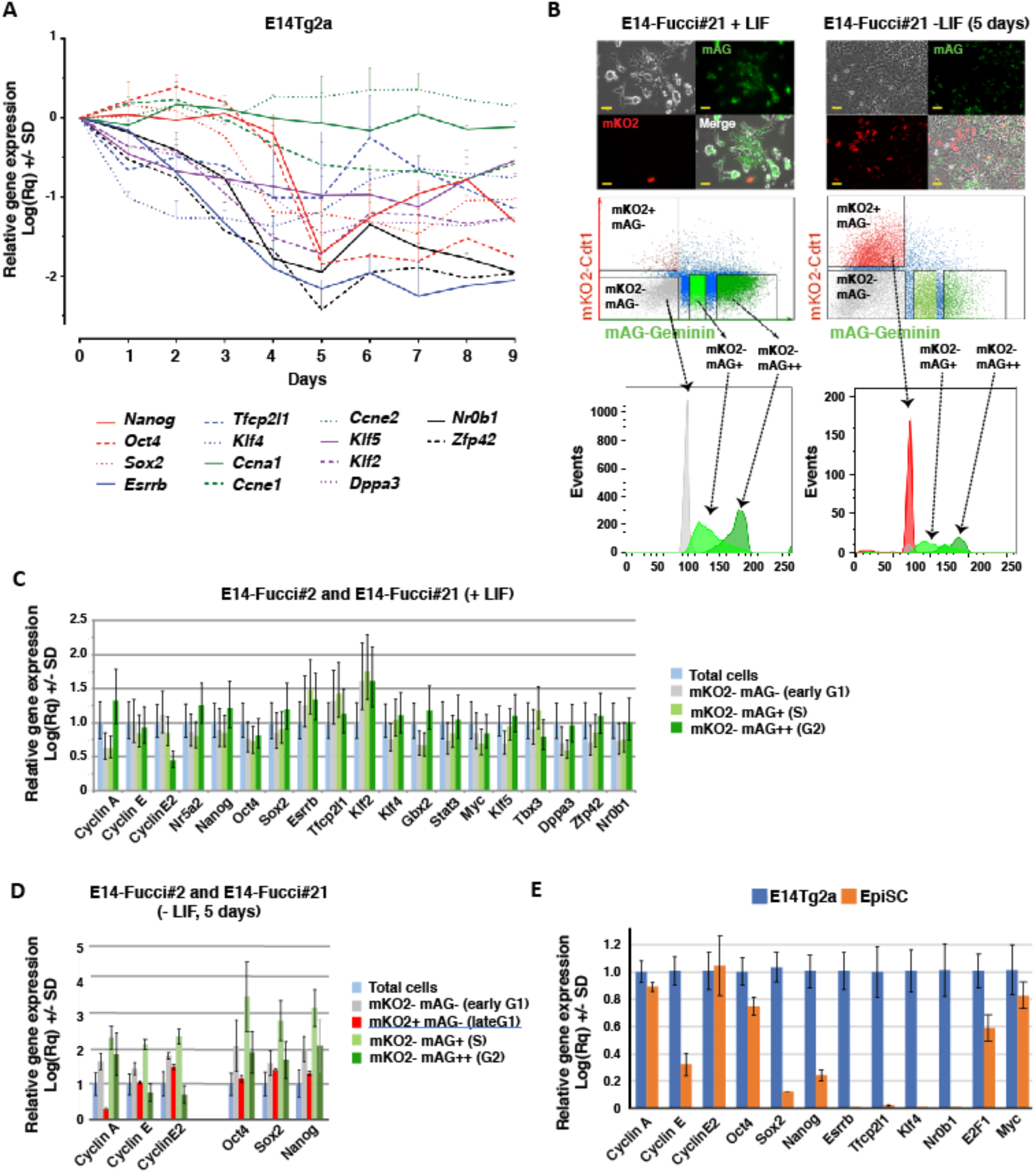
Transcript levels for G_1_ cyclins and pluripotency factors during cell cycle progression. (**A**) Gene expression levels measured by qRT-PCR in E14Tg2a ESCs, before and after differentiation to embryoid bodies, after normalization to β–actin (*Actb*) and levels measured on day 0. (**B**) E14Tg2a–Fucci ESCs, expressing mKO2-hCdt1 and mAG–hGeminin, before and after withdrawal of LIF for 5 days (scale bar represents 20 μM). Top panel: representative fluorescence image. Mid panel: flow cytometry analysis of E14Tg2a–Fucci ESCs showing the distribution of mKO2(−) mAG(−), mKO2(+) mAG(−), mKO2(−) mAG(+) and mKO2(−) mAG(++) cells [19]. Lower panel: cell population histogram of E14Tg2a-Fucci ESCs showing the DNA content of mKO2(−) mAG(−), mKO2(+) mAG(−), mKO2(−) mAG(+) and mKO2(−) mAG(++) cells after propidium iodide staining. (**C**) Gene expression levels measured by qRT-PCR after fluorescence-activated cell sorting (FACS) of E14Tg2a–Fucci ESCs in three distinct fractions corresponding to cells in the G_1_ [mKO2(−) mAG(−)], S [mKO2(−) mAG(+)], and G_2_ [mKO2(−) mAG(++)] phases, respectively. Transcript levels are normalized to β-actin and the level measured in the total population prior to FACS. (**D**) Gene expression levelsmeasured by qRT-PCR after FACS of LIF-deprived E14Tg2a–Fucci cells in four distinct fractions corresponding to cells in the early G_1_ [mKO2(−) mAG(−)], late G_1_ [mKO2(+) mAG(−)], S [mKO2(−) mAG(+)], and G_2_ [mKO2(−) mAG(++)] phases, respectively. Expression levels are normalized to β–actin (*Actb*) and the level measured in the total population prior to FACS. (**E**) Gene expression levels measured by qRT-PCR in E14Tg2a ESCs and EpiSC, normalized to β–actin (*Actb*) and expression measured in E14Tg2a ESCs. (**A, C-E**) Means and standard deviations calculated from three independent experiments are shown.

### Mapping of cis-regulatory elements in the 5’ flanking region of *Cyclin E*

We examined the transcriptional regulation of *Cyclin E* by transcription factors of the naïve pluripotency network. The distribution of DNAse I hypersensitivity sites was analyzed over a 15 kb region encompassing the *Cyclin E* transcription start site in two mouse ESC lines, CJ7 and E14Tg2a, using data available in the mouse ENCODE data base. We identified three regions hypersensitive to DNAse I in the 5’ flanking region: region I between the transcription start site and −1.2 kb, region II between −4.5 and −6 kb, and region III between −9.8 and −11.5 kb (**Fig. 2A**). In both Bruce4 and E14Tg2a ESCs, region I and II displayed a strong and moderate enrichment in H3K4me3 histone marks, respectively. The transcription-promoting activity of the three regions was analyzed using a luciferase assay after transfection into E14Tg2a ESCs (**Fig. 2B**). Region I showed high transcriptional activity, region II enhanced this activity by a factor of two, and region III reduced it by a factor of two. Regions I, II, and III were thereafter called “promoter” (P), “Proximal Enhancer” (PE), and “Distal regulator Element” (DE), respectively. After transfection into a mouse fibroblast (STO) cell line, luciferase activity was dramatically reduced compared with that observed in ESCs, in line with the reduced transcript levels observed after ESC differentiation. Moreover, in contrast to the situation observed in ESCs, the PE region had no enhancer activity on transcription initiated from the P region.

**Figure 2:**
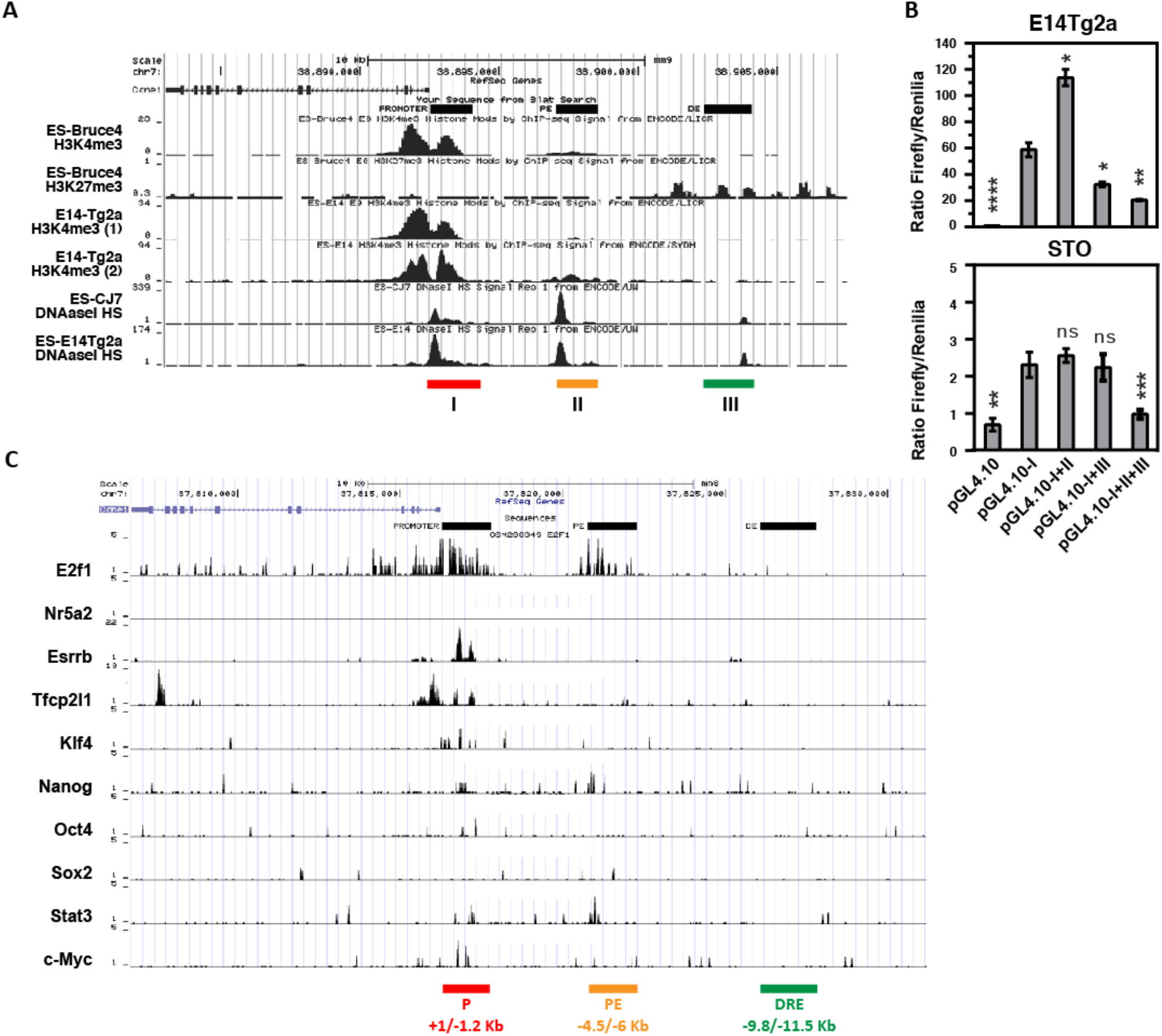
In silico analysis and transcriptional activity of *Cyclin E* promoter region. (**A**) Mapping of DNAse I hypersensitive sites, H3K4me3 and H3K27me3 binding in the 5’ flanking and coding regions of *Cyclin E* identified from ENCODE database [GSM1003830 (DNAseDgf on mESC–CJ7), GSM1014154 (DNAseHS on mESC–E14), and GSM1014187 (DNAseHS on mESC–CJ7), GSM769008 (H3K4me3 on mESC-Bruce4), GSM1000089 (H3K27me3 on mESC-Bruce4), and GSM1000124 (H3K4me3 on mESC-E14) datasets]. (**B**) Luciferase assay to measure the transcriptional activity of regions P (I), PE (II), and DE (III) in E14Tg2a cells (top panel) and STO cells (bottom panel). Mean and standard deviations calculated from three independent experiments are shown and two-way Welch test analysis was used to assess significance (ns p≥0.05, *p<0.05, **p<0.01, ***p < 0.001, ****p < 0.0001). (**C**) Binding of transcription factors to P (I), PE (II), and DE (III) regions of the *Cyclin E* 5’ flanking sequence identified from published previously ChIP-seq data [36, 37] [GSM288349 (E2f1), GSM470523 (Nr5a2), GSM288355 (Esrrb), GSM288350 (Tfcp2l1), GSM288354 (Klf4), GSM288345 (Nanog), GSM288346 (Oct4), GSM288347 (Sox2), GSM288353 (Stat3), GSM288356 (c-Myc), and GSM1208217 (Klf4) datasets].

Using the JASPAR database [39], putative binding sites for the pluripotency regulators Oct4, Sox2, Esrrb, Tfcp2l1, Klf4, Klf5, Tcf3, and STAT3 were identified in the P and PE regions (**Suppl. Fig. 2**). Binding sites for E2F1 and Nr5a2 were also identified in the P region as previously reported [3, 4, 51]. Binding of transcription factors to their respective sites was analyzed from published ChIP-seq data [36, 38]. We identified a strong enrichment in P region-specific sequences after chromatin immunoprecipitation with Esrrb, Tfcp2l1, Klf4, and E2F antibodies, but not with Oct4, Sox2, Nanog, Stat3, Nr5a2 and cMyc antibodies (**Fig. 2C**). Based on these results, all subsequent analyses focused on the role of Esrrb, Tfcp2l1, and Klf4 in the transcriptional regulation of *Cyclin E* via the P region.

### Esrrb, Tfcp2l1, and Klf4 binding sites in the promoter region of *Cyclin E*

The P region contains two binding sites for Klf4 at positions −9/−19 and −185/−195 with respect to transcription start site (relative scores of 98.4% and 95.7%, respectively), one binding site for Tfcp2l1 at position −390/−404 (relative score of 90.2%), and two binding sites for Esrrb at positions −538/−548 and −870/−880 (relative scores of 97.4% and 96.3%, respectively) (**Figure 3A**). The role of the five binding sites in *Cyclin E* transcription was explored by site-directed mutagenesis and analysis of P region transcriptional activity in a luciferase assay (**Figure 3B**). Mutation of Klf4 binding sites had no significant effect on transcriptional activity. Mutation of the Tfcp2l1 and the proximal Esrrb binding sites had a moderate effect on transcription of luciferase (reduction of transcriptional activity to 20% and 58% relative to wild type P region, respectively). In contrast, mutation of the distal Esrrb binding site reduced transcriptional activity to 99% of the wild type sequence (**Figure 3B**). None of these mutations significantly altered the transcriptional activity of the P region when transfected into STO fibroblast cells. We concluded that the distal Esrrb binding site was essential to the transcriptional activity of the promoter element of *Cyclin E* in ESCs, and the proximal Esrrb site and the Tfcp2l1 site play ancillary roles. Luciferase activity associated with pGL4.10-P was also strongly reduced in EpiSC as compared to ES cells (**Figure 3c**), suggesting that the *Cyclin E* promoter region is less active in the primed state than in the naïve state of pluripotency.

**Figure 3:**
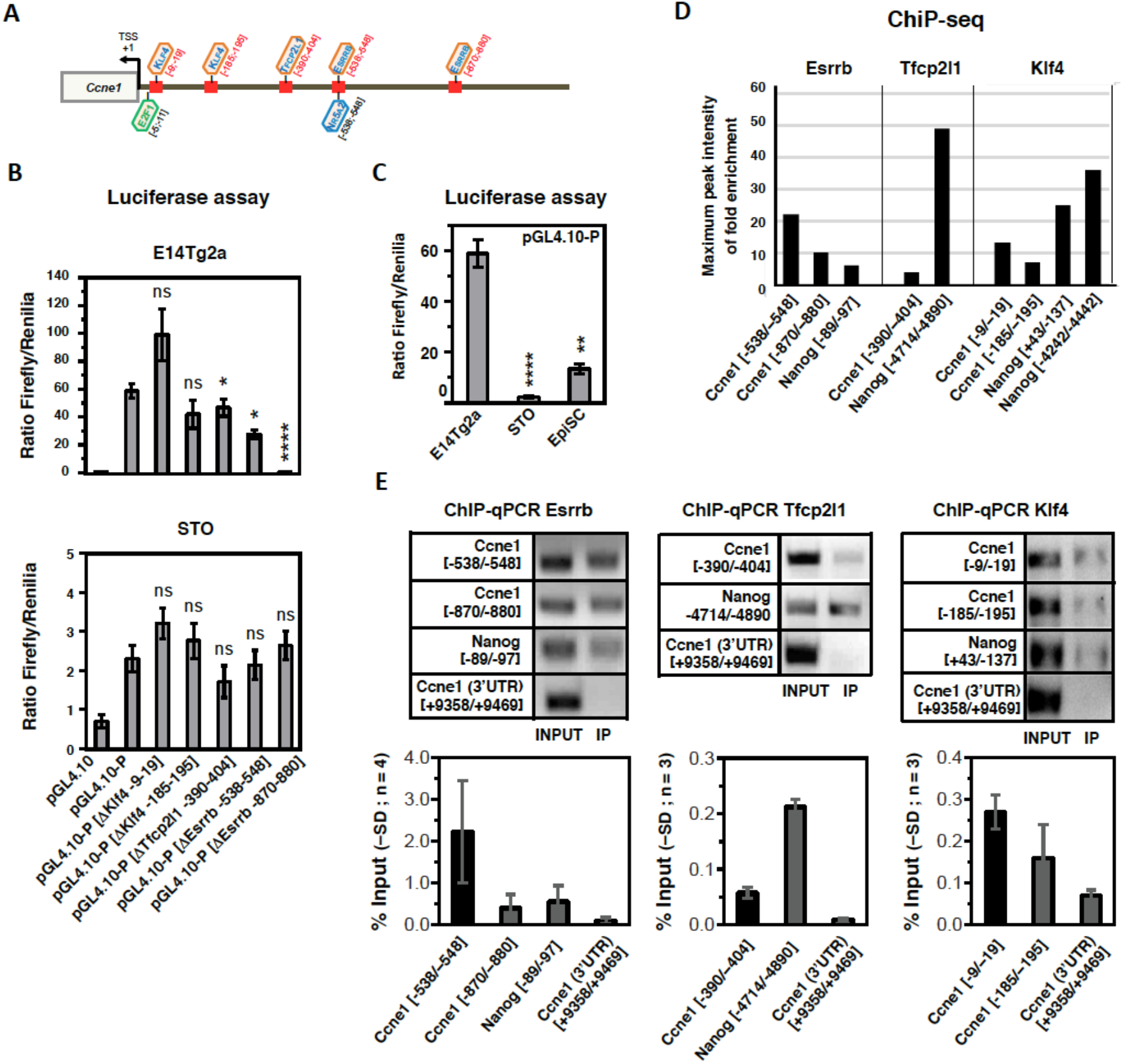
In silico analysis of the P region of the *Cyclin E* gene. (**A**) Mapping of Esrrb, Klf4, Tfcp2l1, E2F, and Nr5a2 putative transcription factor binding sites as determined by Transcription Factor Binding Sites annotation (TFBS) public databases as well as published data. (**B**) Luciferase assay to measure the transcriptional activity of the P region after disruptive mutations of the putative Esrrb, Klf4, and Tfcp2l1 binding sites in E14Tg2a cells (top panel) and STO cells (bottom panel). (**C**) Luciferase assay to measure the transcriptional activity of the P region of Cyclin E after transfection of pGL4.10-P in E14Tg2a cells, EpiSC and STO cells. (**D**) Binding of Esrrb, Klf4, and Tfcp2l1 to the P region of *Cyclin E* and to the promoter of *Nanog* determined by ChIP-seq (data mining from [36] and [38] [GSM288350 (Tfcp2I1), GSM288355 (Esrrb), and GSM1208217 (Klf4) datasets]). Graph represents the maximum enrichment value for each region of interest. (**E**) Binding of Esrrb, Klf4, and Tfcp2l1 to the P region of *Cyclin E* and to the promoter of *Nanog* determined by ChIP–qPCR. (**B,C**) Means and standard deviations calculated from at least three independent experiments are shown and two-way Welch test analysis was used to assess significance (ns p≥0.05, *p<0.05, **p<0.01, ***p < 0.001, ****p < 0.0001).

Binding of Esrrb, Tfcp2l1, and Klf4 to the P region of *Cyclin E* was measured using previously published ChIP-seq data [36]. We observed a high occupancy of Esrrb binding sites at positions −538/−548 and −870/−880 (**Figure 3D; Suppl. Fig. 3**). For comparison, the Esrrb binding site found at position −89/−97 in the *Nanog* promoter [52] showed only a low occupancy. In contrast, we observed a low occupancy of Tfcp2l1 and Klf4 binding sites at position −390/−404, −9/−19 and −185/−195 in the *Cyclin E* promoter, compared with a high occupancy at position −4714/−4890, +43/−137 and −4242/4442 in the *Nanog* promoter [34, 53]. These data indicate a strong interaction of Esrrb and a much weaker interaction of Tfcp2l1 and Klf4 to their respective predicted binding sites in the *Cyclin E* promoter. Importantly, these results could be corroborated by ChIP-qPCR (**Figure 3E**). Strong enrichment in PCR fragments encompassing the Esrrb binding sites in the *Cyclin E* promoter were observed compared with moderate enrichment in fragments encompassing the Klf4 binding sites and weak enrichment in a fragment encompassing the Tfcp2l1 binding site. As the two Esrrb binding sites are located in close proximity, we could not assess whether Esrrb shows a similar affinity to the proximal as well as the distal predicted binding site. Altogether, these results indicate a significant regulatory role of Esrrb to the *Cyclin E* promoter, in line with the results of the transcriptional activity of mutant promoters.

### Regulation of *Cyclin E* by Esrrb, Tfcp2l1, and Klf4

To further substantiate the implication of Esrrb, Tfcp2l1 and Klf4 in the transcriptional regulation of *Cyclin E*, their expression was knocked down by means of doxycyclin-induced expression of two independent shRNAs (*Sh#1* and *Sh#2*) in KH2 ESCs [48]. The expression levels of *Esrrb, Klf4*, and *Tfcp2l1* could be reduced to less than 10% after 48 h induction with doxycycline for sh#1 (**Figure 4A**). *Cyclin E* transcript levels showed a correlated decrease compared to control cells after knockdown of *Esrrb* (77%), *Tfcp2l1* (69%) and *Klf4* (74%), respectively. In ESCs expressing *Sh#2*, doxycycline treatment resulted in a substantially smaller reduction of *Esrrb, Tfcp2l1*, and *Klf4* transcript levels than Sh#1 (i.e. to 25%, 63%, and 30% of their original levels, respectively). Nevertheless, the *Cyclin E* transcript level was significantly reduced (65%, 61%, and 24%, respectively). No alteration of *Esrrb, Tfcp2l1, Klf4*, or *Cyclin E* transcript levels was observed in control cells expressing *Sh-Control*. In accordance with the reduced transcripts, Cyclin E protein levels decreased to 25% and 50% of the level measured in control cells after knockdown of *Esrrb* and *Tfcp2l1*, respectively (**Figure 4B**). ESCs expressing *Sh-Esrrb* and *Sh-Tfcp2l1* showed only minor alterations in the expression of other pluripotency markers including *Nanog, Oct4, Sox2, Klf2, Klf5, Tbx3, Nr0b1*, and *Zfp42*, indicating that the observed reduction of *Cyclin E* transcript levels after *Esrrb* and *Tfcp2l1* knockdown is not a consequence of differentiation (**Suppl. Fig. 4**). In contrast, *Klf4* knockdown resulted in a significant attenuation of most of the aforementioned pluripotency regulators. As the Klf4 binding sites are apparently not involved in the transcriptional regulation of *Cyclin E*, the observed downregulation of *Cyclin E* in *shKlf4* ESCs might be explained by a substantial rate of spontaneous differentiation. In the next step, Esrrb, Klf4, and Tfcp2l1 were overexpressed using a doxycycline-inducible vector system in KH2 ESCs (**Figure 4C**). This resulted in a 2.7- and 2.2-fold increase in *Cyclin E* transcript levels after Esrrb and Klf4 overexpression, respectively. No significant increase was observed after Tfcp2l1 overexpression. These results were confirmed by the analysis of Cyclin E, Esrrb, Tfcp2l1, and Klf4 protein levels (**Figure 4D**). For Tfcp2l1 overexpression, Cyclin E increase was observed only at the protein level, which may suggest a post-transcriptional regulation. Together, these results indicate a strong correlation between the expression levels of Esrrb and Cyclin E.

**Figure 4:**
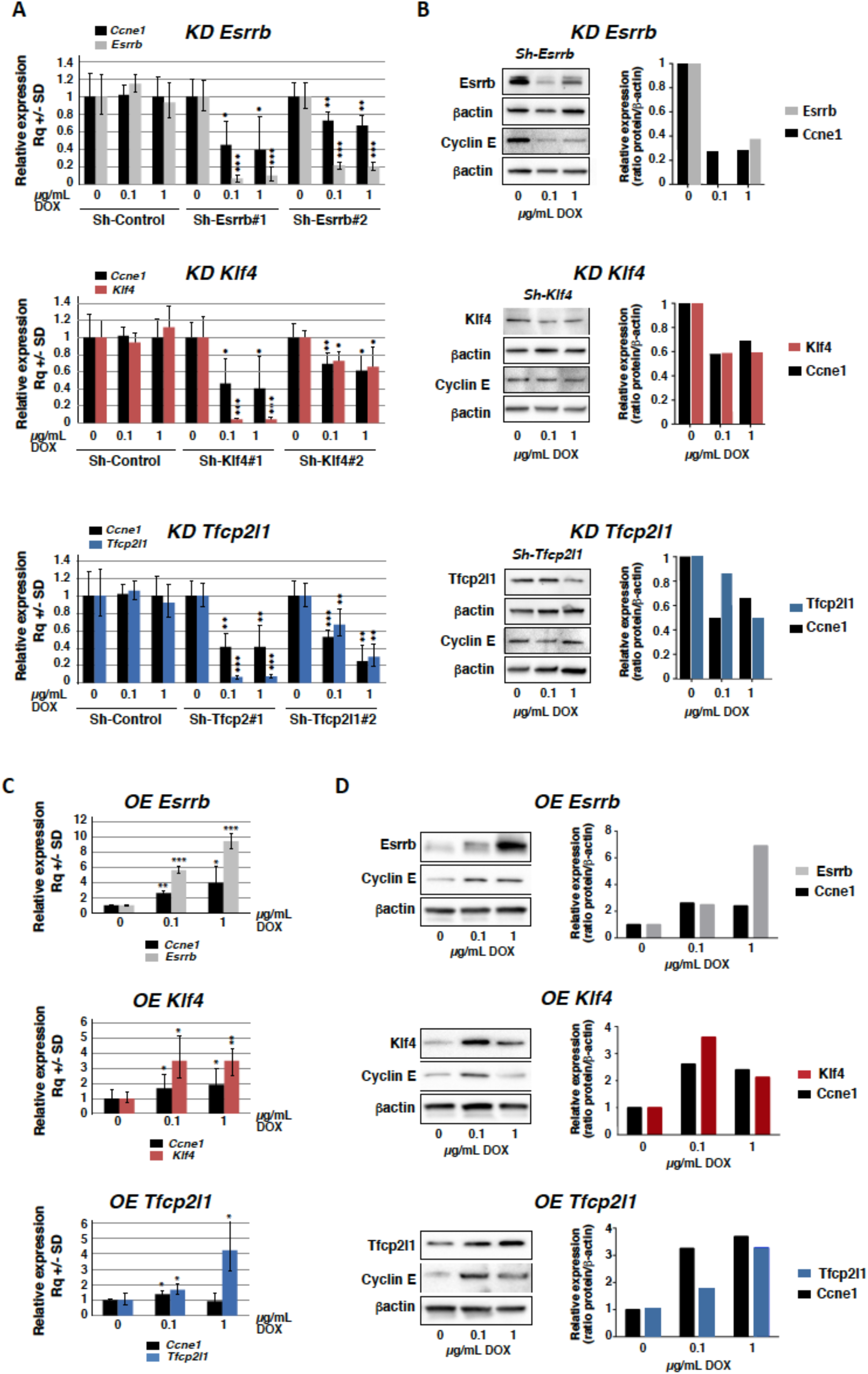
Regulation of *Cyclin E* expression by Esrrb, Klf4, and Tfcp2l1. (**A**) Doxycycline-induced expression of sh-Esrrb#1, sh-Esrrb#2, sh-Klf4#1, sh-Klf4#2, sh-Tfcp2l1#1, sh-Tfcp2l1#2, and sh-Control in KH2 ESCs. Expression of the indicated genes is measured by qRT-PCR after 48 h of treatment with doxycycline at 0.1 and 1.0 μg/mL. All gene expression values are normalized to the value measured in the absence of doxycycline (0 μg/mL). (**B**) Doxycycline-induced expression of sh-Esrrb#1, sh-Klf4#1, and sh-Tfcp2l1#1 in KH2 ESCs. Expression of the indicated genes was detected by immunoblotting after 48 h of treatment with doxycycline at 0.1 and 1.0 μg/mL and compared with the absence of doxycycline (0 μg/mL). (**C,D**) Doxycycline-induced expression of Esrrb, Klf4, and Tfcp2l1 mouse cDNAs in KH2 ESCs. Expression of the indicated genes was measured by qRT-PCR (**C**) and immunoblotting (**D**) and after 48 h of treatment with doxycycline at 0.1 and 1.0 μg/mL, and compared with the absence of doxycycline (0 μg/mL).). (A,C) Means and standard deviations (SD) calculated from three independent experiments are shown and twoway Welch test analysis was used to assess significance (ns p≥0.05, *p<0.05, **p<0.01, ***p < 0.001, ****p < 0.0001).

### Partial rescue of Esrrb knockdown-induced differentiation by Cyclin E

Several studies have pointed to Esrrb as an inducer of somatic cell reprogramming and ESC self-renewal [29, 30, 54]. The role of Esrrb in the regulation of Cyclin E therefore prompted us to investigate the capacity of cyclin E to oppose Esrrb knockdown-induced differentiation. We therefore used EKOiE cells expressing doxycycline-regulated *Esrrb* cDNA in an *Esrrb*-null background [49]. After doxycycline deprivation for 48 h, Esrrb was undetectable. Consequently, a twofold reduction of both *Cyclin E* mRNA and cyclin E protein was observed (**Figure 5A**). Expression of *Tfcp2l1* and *Klf4* were also reduced, which may have contributed to the downregulation of *Cyclin E* (**Figure 5B**). When doxycycline-deprived EKOiE cells were supplemented with doxycycline for 48 h, they restored expression of *Esrrb, Tfcp2l1, Klf4*, and *Cyclin E* to their original levels. Expression of *Oct4* and *Sox2* remained unchanged in this experimental setting. Next, EKOiE cells were transfected with a plasmid carrying a Myc-tagged rat cyclin E cDNA, or with an empty plasmid [55] (**Figure 5C**). EKOiE–^Myc^cyclin E and EKOiE-control cell populations were analyzed using a colony-forming assay to assess the balance between self-renewal and differentiation (**Figure 5D**). Withdrawal of doxycycline resulted in a gradual increase of the proportion of mixed and differentiated colonies indicating that the loss of Esrrb disrupted self-renewal as previously reported [30]. In the presence of doxycyline, no difference was observed between EKOiE–^Myc^ cyclin E and control cells regarding the proportion of undifferentiated, mixed, and differentiated colonies, suggesting that enforced expression of cyclin E has no observable effect on self-renewal in the presence of Esrrb. In contrast, the withdrawal of doxycycline for 3,5, and 7 days, leads to a significantly increased proportion of mixed and undifferentiated colonies in EKOiE–^Myc^ cyclin E cells when compared to control cells. These results strongly suggest that enforced expression of cyclin E opposes ESC differentiation induced by downregulation of Esrrb.

**Figure 5:**
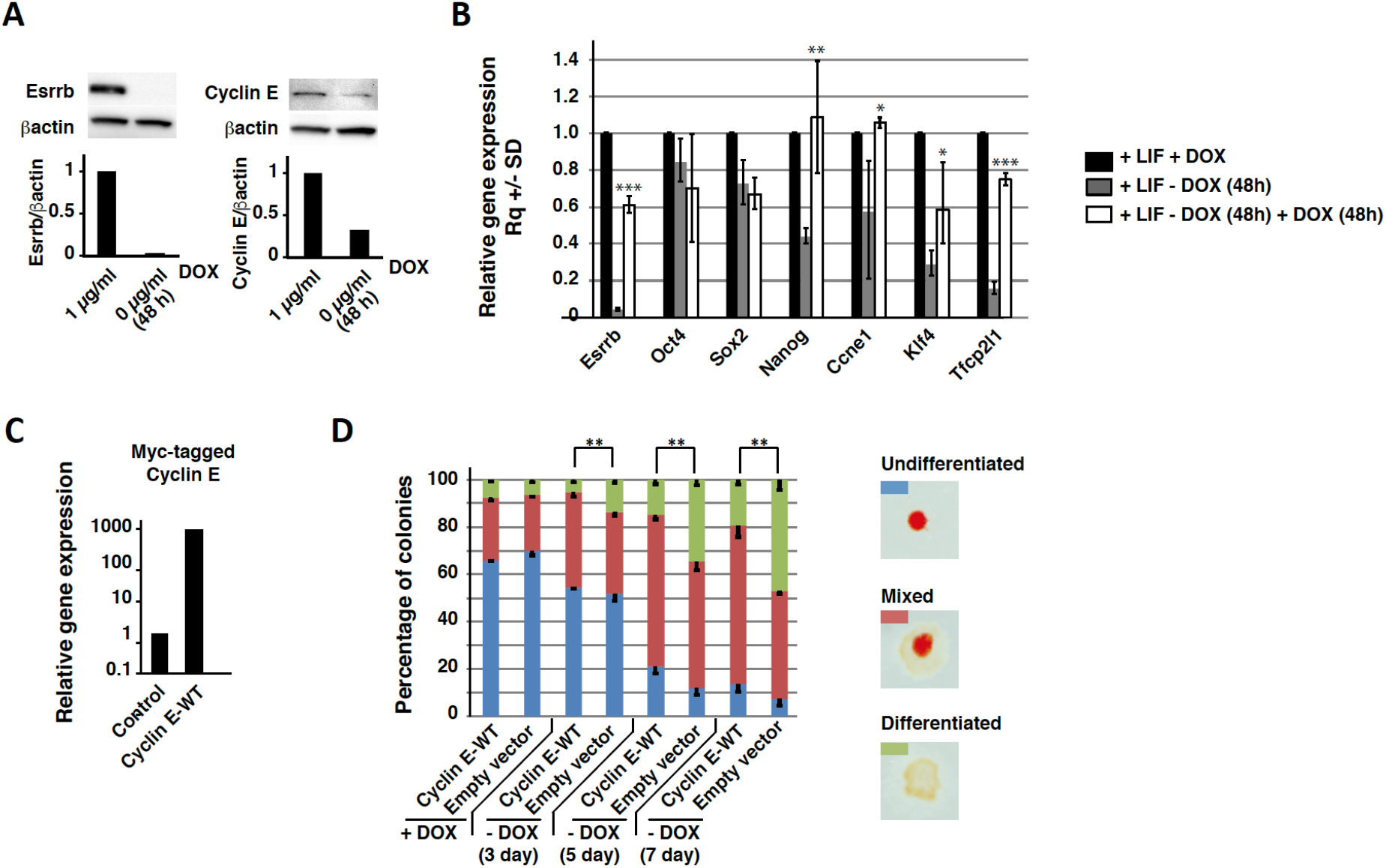
Rescue of Esrrb knockdown-induced differentiation by Cyclin E. (**A**) Expression of Esrrb and Cyclin E in EKOiE cells, before and after withdrawal of doxycycline for 48 h, and analyzed by immunoblotting. Histograms represent the quantification of Esrrb and Cyclin E after normalization to β–actin and to expression in + DOX condition. (**B**) Expression of *Esrrb, Oct4, Sox2, Nanog, Cyclin E (Ccne1), Klf4*, and *Tfcp2l1* in EKOiE cells (+ LIF + DOX), after withdrawal of doxycycline for 48 h [+ LIF-DOX (48 h)], and after addition of doxycycline for 48 h [+ LIF-DOX (48 h) + DOX (48 h)], normalized to expression measured in control cells [+ LIF + DOX]. *p < 0.05; **p < 0.01. ***p < 0.001 (three independent experiments). (**C**) qRT-PCR analysis of *Cyclin E* mRNA levels in EKOiE–^Myc^ cyclin E and control EKOiE cells using myc-tagged *Cyclin E*-specific primers. (**D**) Histogram for the proportion of undifferentiated (U), mixed (M), and differentiated (D) colonies in EKOiE cells expressing ^Myc^cyclin E and control vector cells before (+ DOX) and after withdrawal of doxycycline (− DOX) for 3, 5 and 7 days. **p < 0.01 (three independent experiments). Right panel: representative pictures of undifferentiated (U), mixed (M), and differentiated (D) colonies after staining for stem cell-specific alkaline phosphatase activity.

### Regulation of *Cyclin E* transcript level by miR-15a

MicroRNA miR-15a was shown to act as a negative regulator of *Cyclin E* in somatic cells [7, 8]. We therefore asked if the decrease in *Cyclin E* transcript levels observed during ES cell differentiation could also be a result of a rise in *miR-15a* levels. A qRT-PCR analysis revealed that changes in miR-15a expression mirrored those of *Cyclin E* between 1 and 9 days of differentiation, suggesting a cross-regulation (**Fig. 6A**). To demonstrate the regulation of *Cyclin E* expression levels by miR-15a, E14Tg2a ESCs were transfected with a miR-15a-5p mimic, resulting in a 58% reduction of *Cyclin E* transcripts levels after 48 hours of culture. Nanog transcript levels were also decreased, albeit to a lesser extend (41%) (**Fig. 6B**). These results strongly suggest that the reduced levels of Cyclin E upon differentiation might at least partially be attributed to an increased expression of miR-15a.

**Figure 6:**
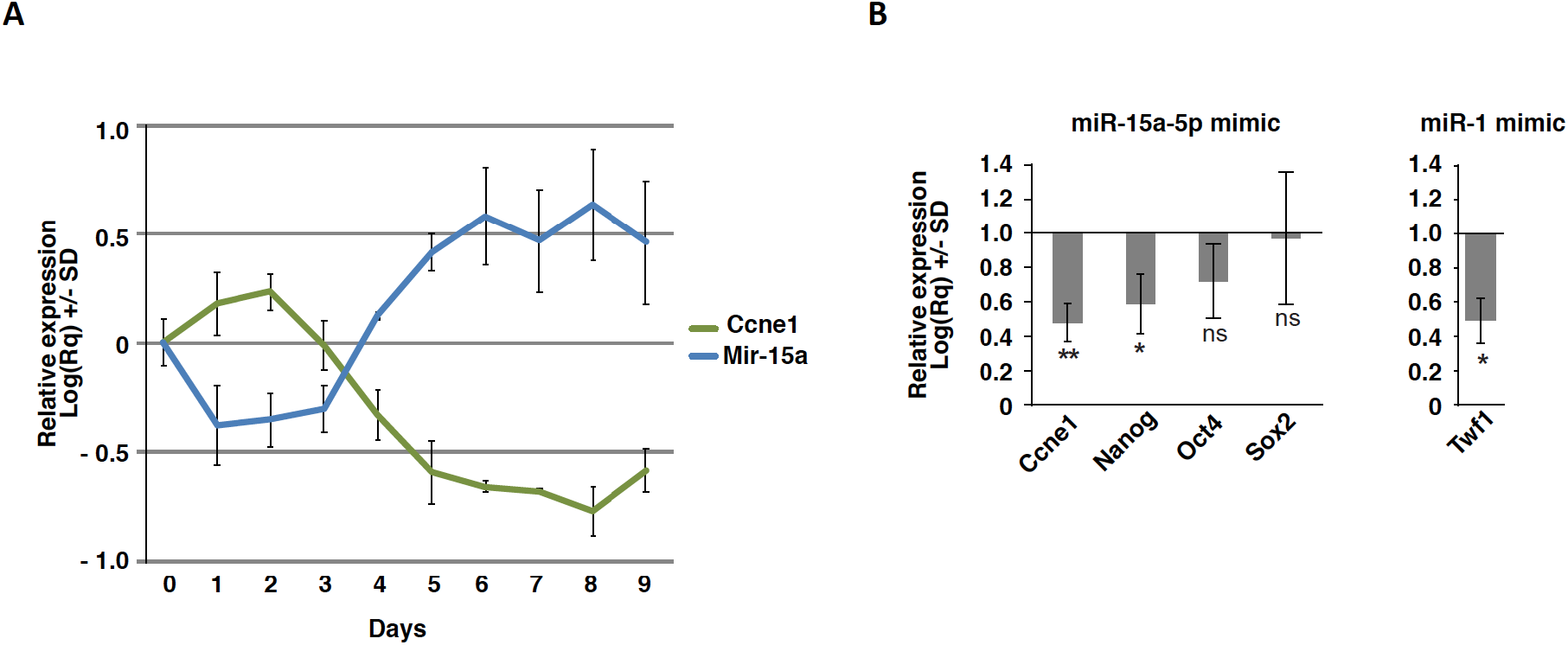
Regulation of *Cyclin E* transcript level by miR-15. (**A**) RNA expression levels of *Cyclin E* and miR-15a measured by qRT-PCR in E14Tg2a before and after differentiation to embryoid bodies (1-9 days), after normalization to β-actin (*Actb*) and levels measured on day 0. (**B**) *Cyclin E* (*Ccne1*), *Nanog, Oct4* and *Sox2* transcript levels measured by qRT-PCR 48 hours after transfection of 25 mM miR-15a-5p mimic in E14Tg2a ESCs, normalized to β–actin (*Actb*) and levels measured on untransfected cells. Transfection of a miR-1 mimic and qRT-PCR analysis of *Twf1* transcripts is used as control. Three independent experiments are shown and two-way Welch test analysis was used to assess significance (ns p≥0.05, *p<0.05, **p<0.01).

## Discussion

In somatic cells, expression of cyclin E varies during the cell cycle, reaching a maximum at the G_1_/S phase boundary. Repression of the *cyclin E* gene during G_2_/M and the early G_1_ phase of the cell cycle are mediated through the assembly of a multiprotein complex containing hypophosphorylated Rb, a histone deacetylase as well as the SWI/SNF chromatin remodeling complex, which bind to the *cyclin E* promoter in order to silence transcription. Transcriptional activation of the *cyclin E* gene during progression through the G_1_ phase depends on the activity of cyclin D/Cdk4 and cyclin D/Cdk6 complexes, which phosphorylate and inactivate retinoblastoma protein (pRb) leading to the release of the repressor proteins of the E2F-transcription factor family [56]. In mouse ESCs, cell cycle regulation of *Cyclin E* seems to obey a different rule. Despite very low levels of D-type cyclins [12, 17] [57], the G_1_-specific hypophosphorylated form of Rb is almost absent [17, 58], and essentially all E2F transcription factors are free from Rb proteins and bind the cyclin E promoter to stimulate transcription, regardless of the cell’s growth cycle position [18]. The theory of a continuous and uniform expression of *Cyclin E* was challenged in a recent study using ESCs synchronized with nocodazole, showing that *Cyclin E* transcripts indeed reached a maximum during G_1_ phase [59]. Strikingly, a similar cell cycle-dependent pattern was observed for Nanog and Esrrb. In sharp contrast, we did not observe a similar pattern using nonsynchronized ESC–Fucci, which raises the question of whether chemically-synchronized ESC can regulate gene expression after release from the mitotic block.

Esrrb, Tfcp2l1, and Klf4 are three transcription factors implicated in the control of naive pluripotency. Esrrb is a direct target gene of both Nanog and the GSK3/β-catenin/Tcf3 pathway [29, 30], and both Tfcp2l1 and Klf4 are direct target genes of the LIF/STAT3 signaling pathway [23, 25, 33]. In the present study, we showed that *Cyclin E* is a direct target of these three transcription factors, pointing to regulation of cyclin E expression by the gene regulatory circuitry that controls naive pluripotency. Among the three factors studied, Esrrb seems to play a major role as shown, first, by the dramatic reduction of transcriptional activity from the promoter region lacking the distal Esrrb binding site, and second, by the downregulation of *Cyclin E* after Esrrb expression has been knocked down or turned off. Moreover, Esrrb overexpression resulted in a 4-fold increase of the steady-state level of *Cyclin E* RNA, further supporting the link between Esrrb and the regulation of *Cyclin E* expression. Tfcp2l1 contributes to the transcriptional regulation of *Cyclin E* in conjunction with Esrrb. The proximal Esrrb binding site overlaps with the binding site for Nr5a2 (LRH-1), a transcription factor involved in the control of naive pluripotency [60]. In addition, Nr5a2 has been shown to regulate expression of *Cyclin E* in conjunction with β-catenin in intestinal crypt cells [51]. In ESCs, mutation of the Esrrb/Nr5a2 binding site has only a minor effect on the transcriptional activity of the promoter region and ChIP-seq studies revealed no binding of Nr5a2 to the promoter region of *Cyclin E*, strongly suggesting that Nr5a2 plays no role in *Cyclin E* transcriptional regulation in ESCs.

We showed that microRNA miR-15a is a negative regulator of Cyclin E in ESCs, in line with its function in somatic cells [7, 8]. Thus, we propose a regulatory model of *Cyclin E* expression, in which the elevated level of *Cyclin E* transcripts observed in ESCs results from both a transcriptional activation by Esrrb and a lack of negative regulation by miR-15a. Differentiation would trigger both the down-regulation of Esrrb and the elevation of miR-15a, resulting in a rapid drop in *Cyclin E* transcript level. Interestingly, miR-15a is a key direct transcriptional target of E2F in somatic cells [8]. Low expression of miR-15a in ESCs and its up-regulation during differentiation strongly suggests that E2F activity is very low in ESCs, and it is only restored after exit from naïve pluripotency.

We showed that mouse EpiSCs, which epitomize the primed state of pluripotency, express *Cyclin E* at a much lower level as compared to the naïve ESCs. We know little about cell cycle regulation in mouse EpiSCs. However, human ESCs, i.e. the human counterparts of mouse EpiSCs, seem to exhibit a somatic-like cell-cycle regulation. In particular, it was observed that *Cyclin E* mRNA levels increases sharply at the G_1_/S transition and that the regulation of *Cyclin E* mRNA expression levels involves the activation of MEK/ERK pathway and the transcription factors c-Myc and E2F [61]. These observations suggest that the transition from naïve-to primed-state pluripotency is accompanied by the loss of the regulation by transcription factors of the naïve pluripotency network and the gain of the regulation by c-Myc and E2F.

## Acknowledgement

This work was supported by the Infrastructure Nationale en Biologie et Santé INGESTEM (ANR-11-INBS-0009), the IHU-B CESAME (ANR-10-IBHU-003), and the LabEx “DEVweCAN” (ANR-10-LABX-0061) and the LabEx “CORTEX” (ANR-11-LABX-0042) of the University of Lyon within the program ‘Investissements d’Avenir’ (ANR-11-IDEX-0007).

---

**Supplementary Figure 1:**
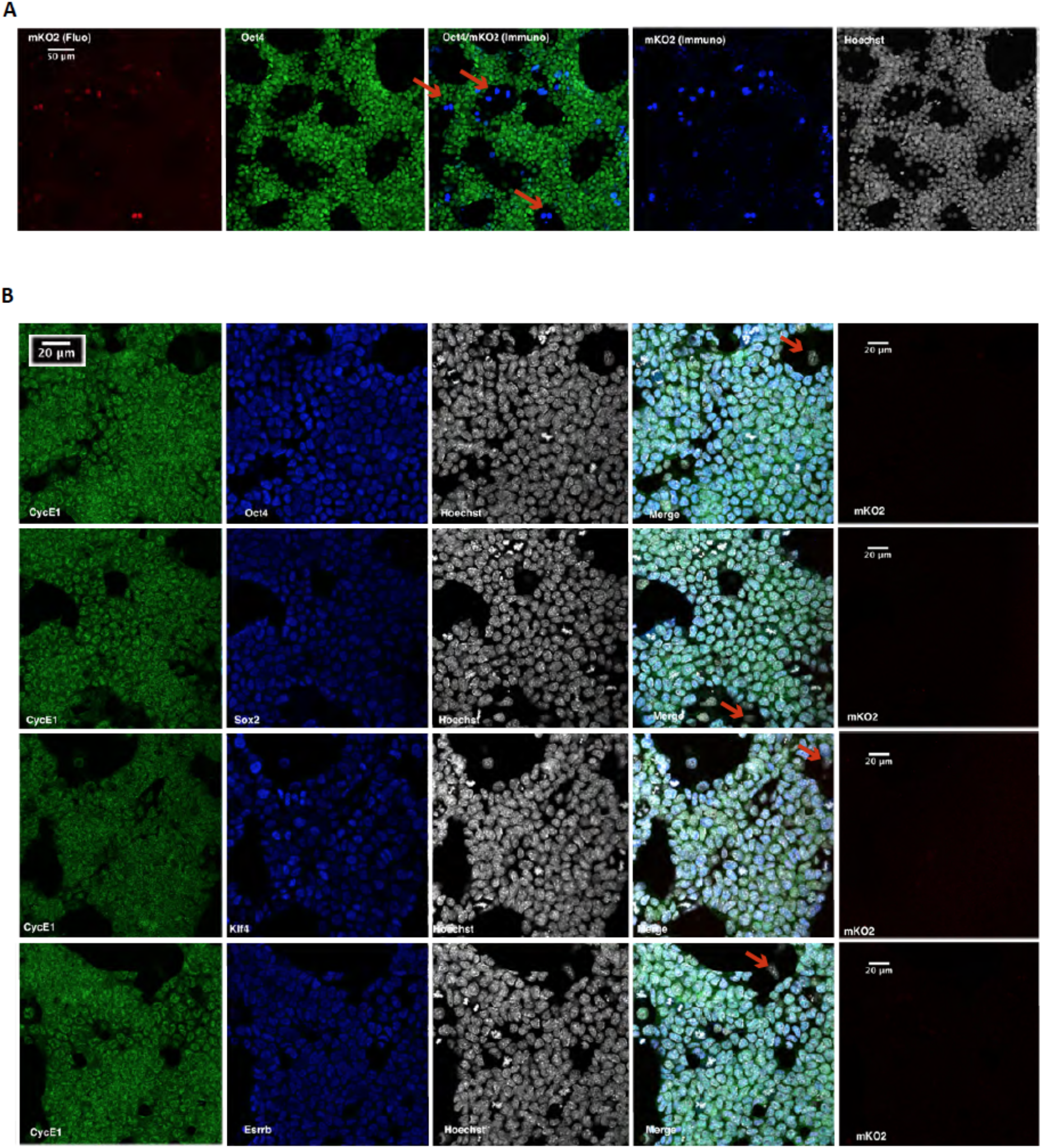
Fluorescence analysis and immunolabeling of E14Tg2a ESC expressing mKO2–hCdt1. (**A**) Representative fluorescence image of E14–mKO2–hCdt1 (red), and immunolabeling for Oct4 (green) and mKO2 (blue). Red arrows indicate mKO2-positive/Oct4-negative cells. (**B**) Immunolabeling of E14–mKO2–hCdt1 cells for cyclin E (CycE1), Oct4, Sox2, Klf4, and Esrrb.

**Supplementary Figure 2:**
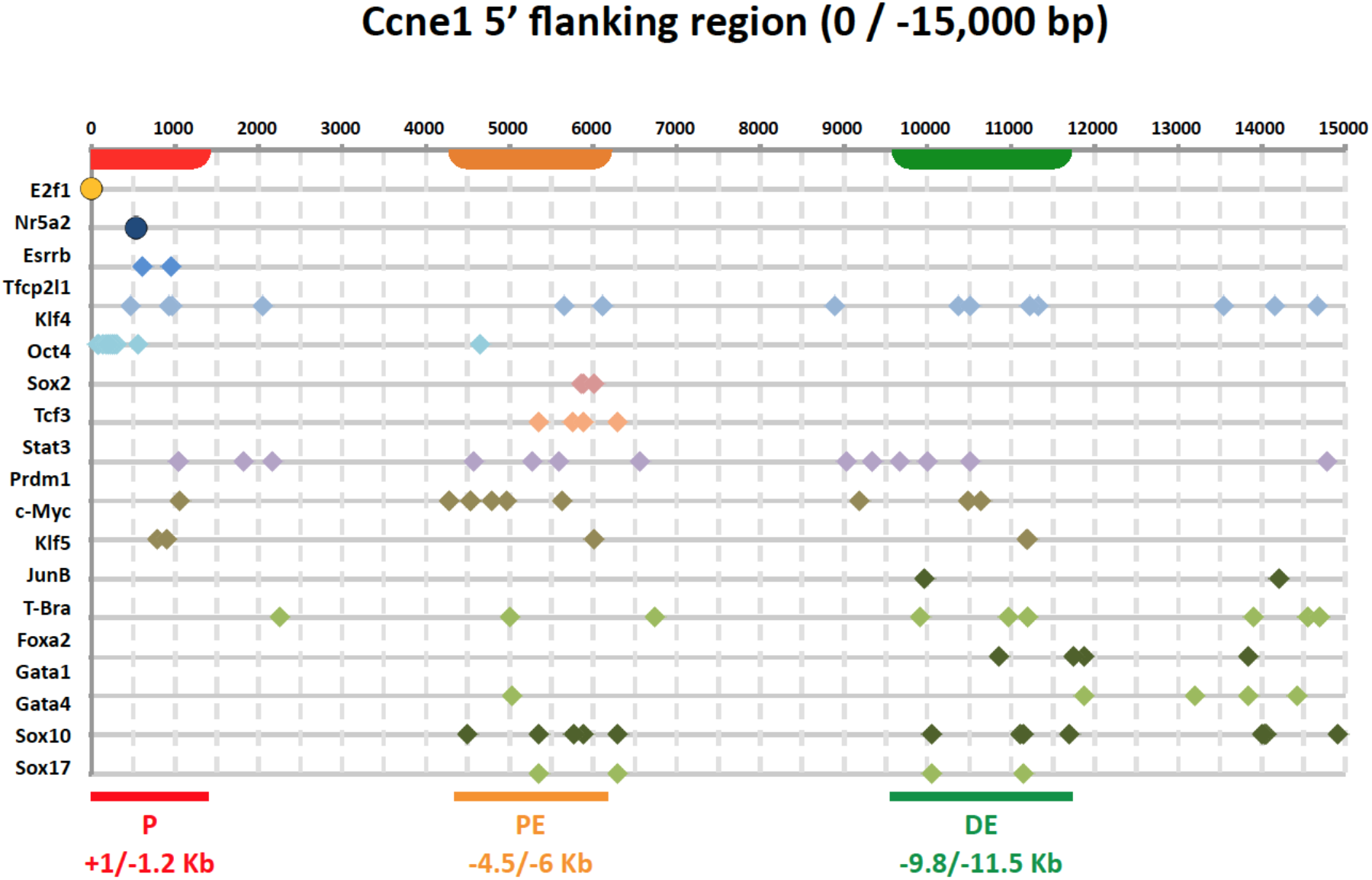
Supplementary Figure 2: Characterization of the *Cyclin E* 5’ flanking sequence. Mapping of putative transcription factor binding sites in the P (I), PE (II), and DE (III) regions of *Cyclin E* 5’ flanking sequence as identified from TFBS public databases.

**Supplementary Figure 3:**
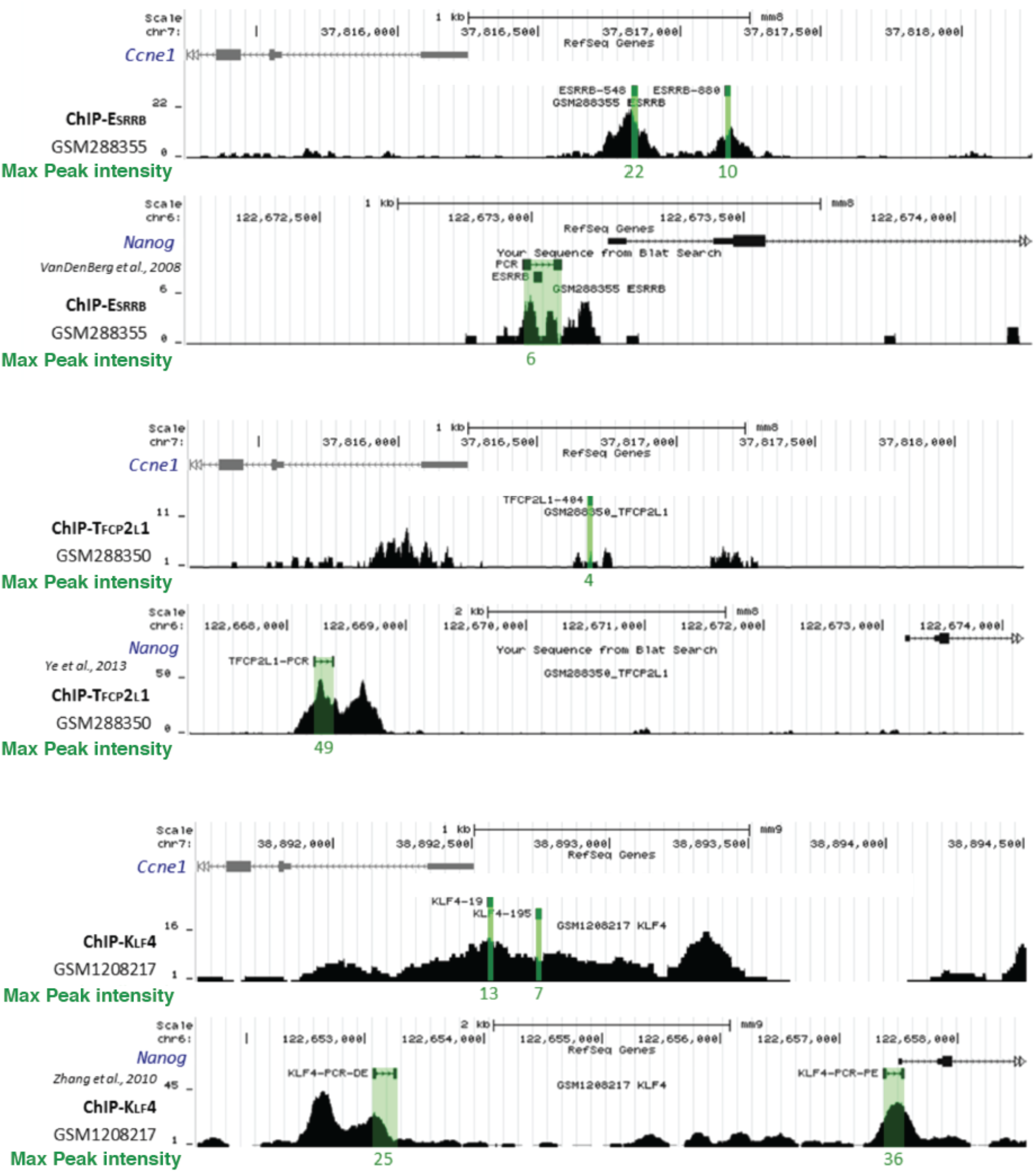
ChIP-seq peaks for Esrrb, Klf4 and Tfcp2l1 for the *Cyclin E* and *Nanog* genes 5’ flanking sequences. Binding of Esrrb, Klf4, and Tfcp2l1 to the P region of *Cyclin E* and to the promoter of *Nanog* determined by ChIP-seq (data mining from [36] and [38] [GSM288355 (Esrrb), GSM288350 (Tfcp2I1), and GSM1208217 (Klf4) datasets]). Putative binding sites on the *Cyclin E* 5’ flanking region and binding-sites, which have already been described on the *Nanog* promoter [Esrrb (VanDenBerg, Mol. Cell. Biol. 2008), Tfcp2l1 (Ye, EMBO J. 2013), and Klf4 (Zhang JBC 2010)] are highlighted.

**Supplementary Figure 4:**
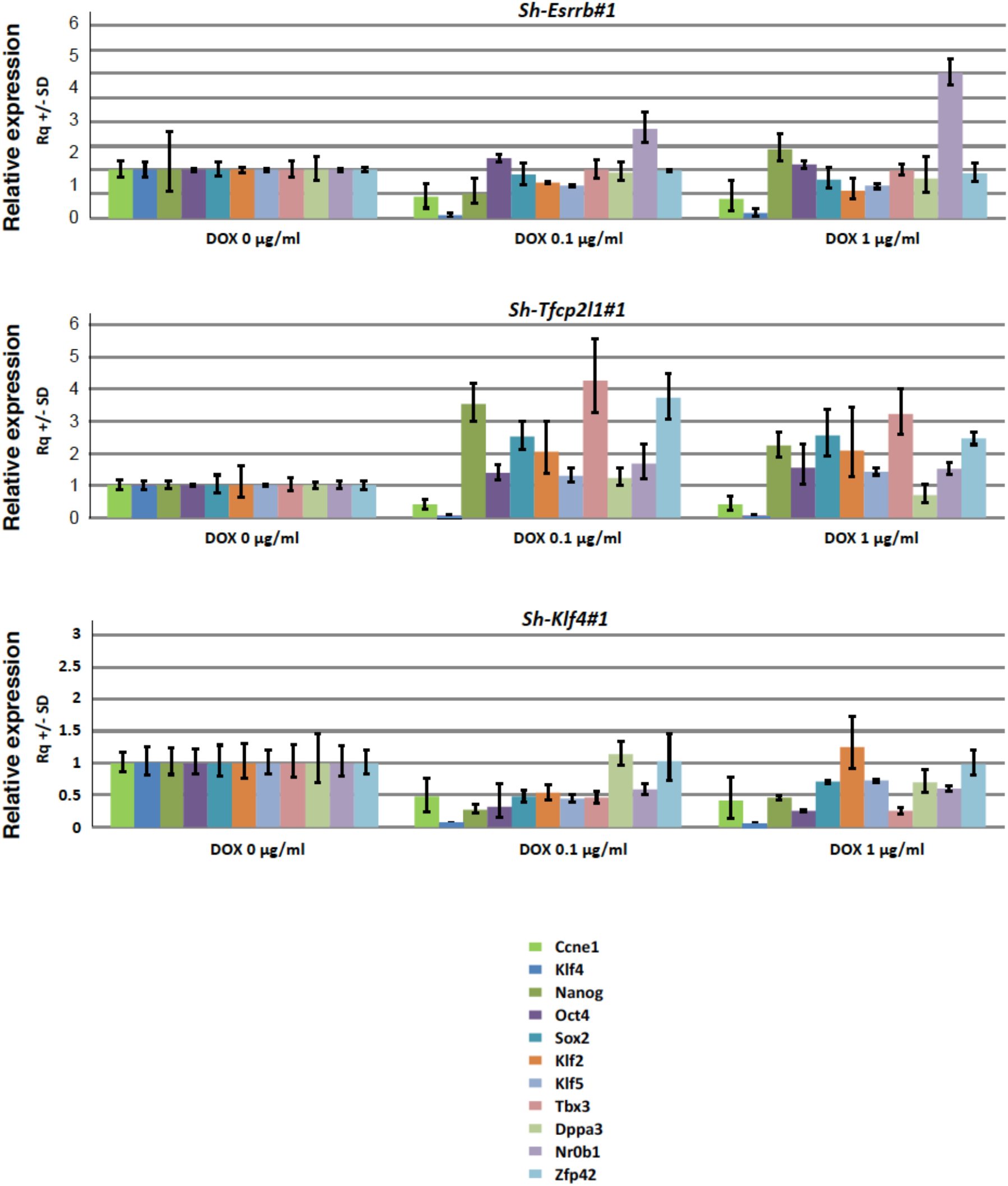
Characterization of KH2 ESCs expressing sh-Esrrb#1, sh-Klf4#1, and sh-Tfcp2l1#1. Doxycycline-induced expression of sh-Esrrb#1, sh-Klf4#1, sh-Tfcp2l1#1, and sh-Control in KH2 ESCs. Expression of the indicated genes is measured by qRT-PCR after 48 h of treatment with doxycycline at 0.1 and 1.0 μg/mL. All gene expression values are normalized to the value measured in the absence of doxycycline (0 μg/mL).

**Table S1:**
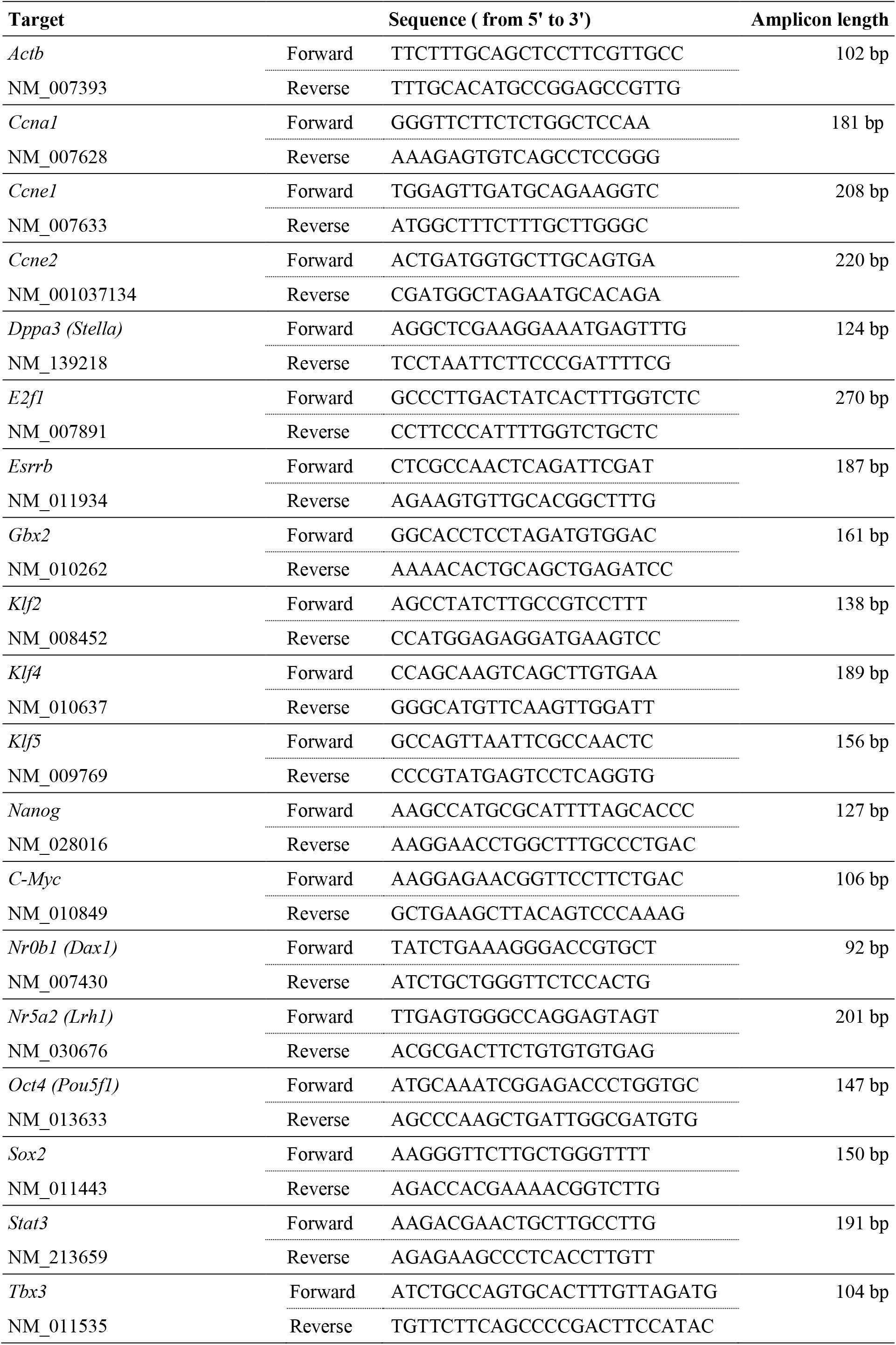
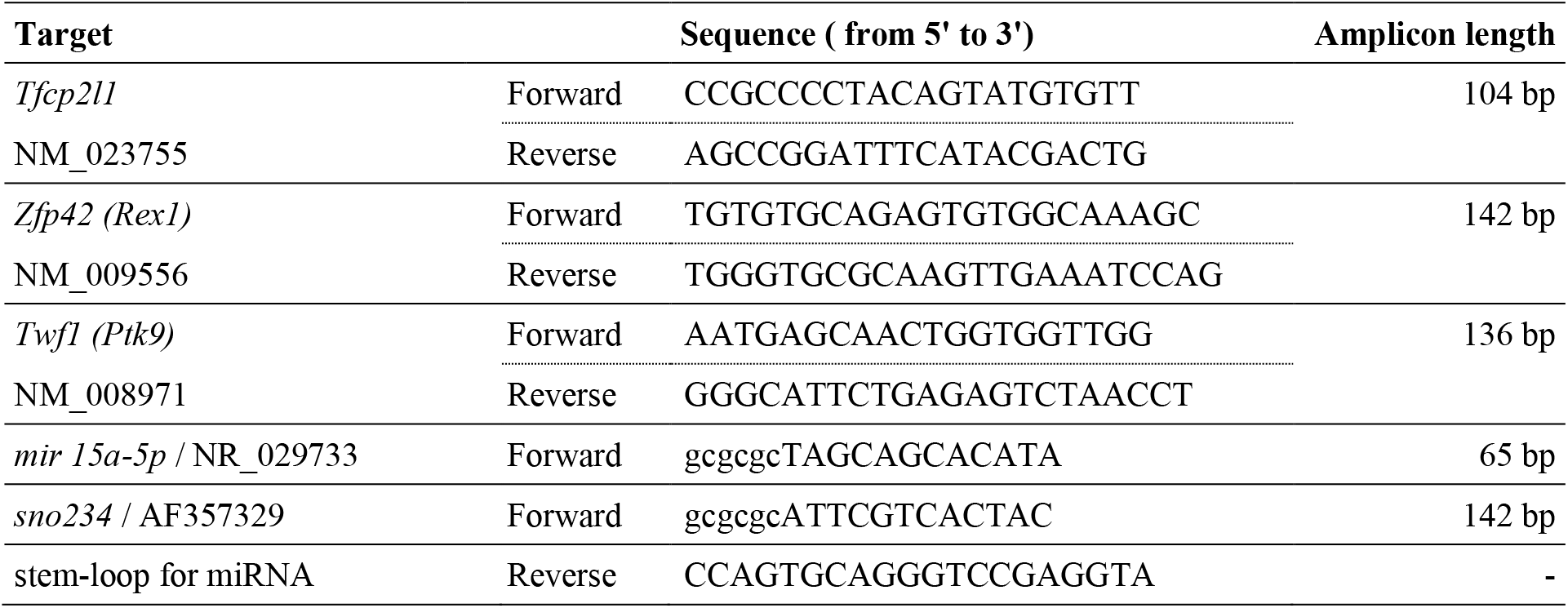
PCR primers for qRT-PCR.

**Table S2:**
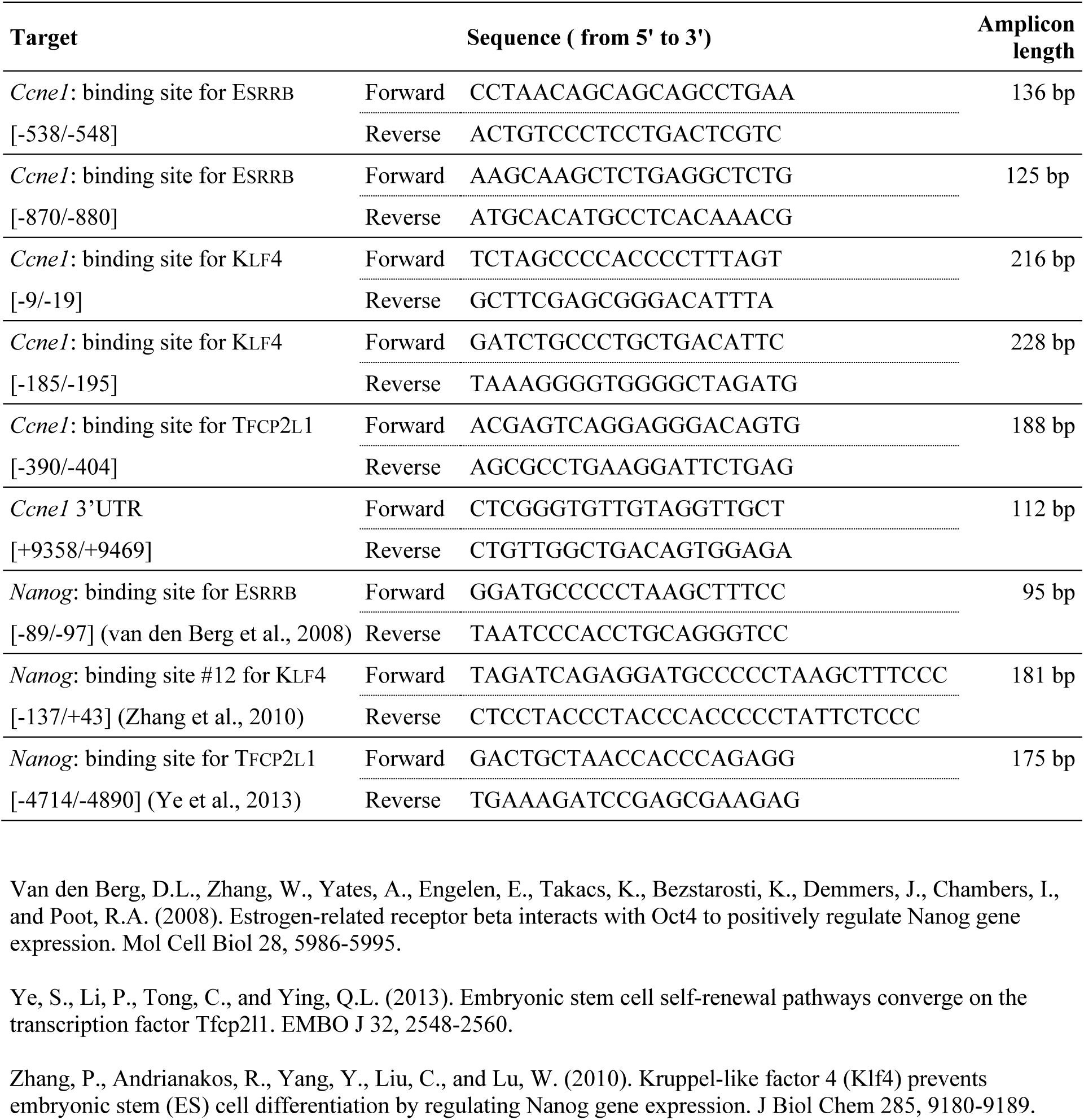
PCR primers for amplification of ChIP products.

**Table S3:**
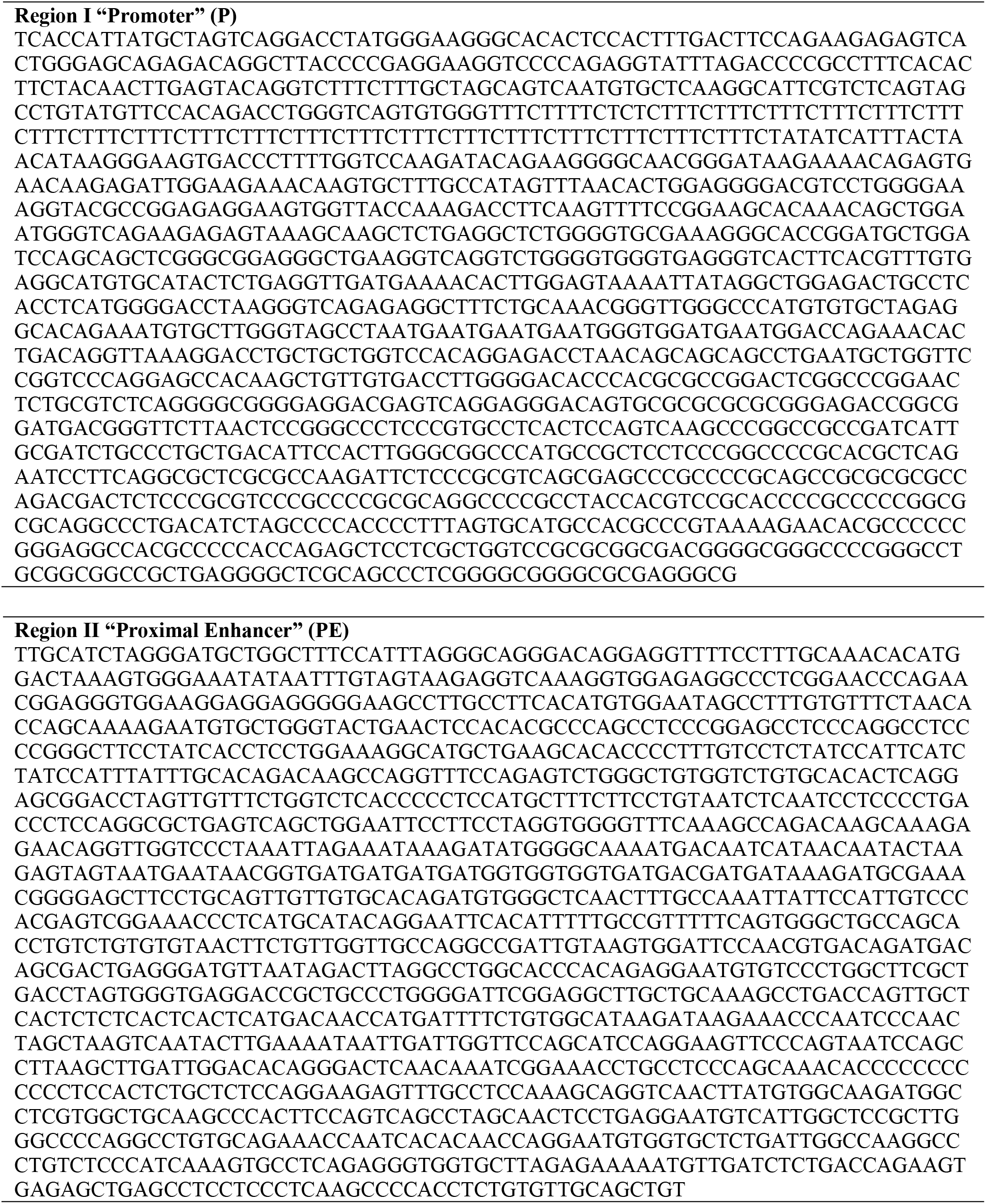
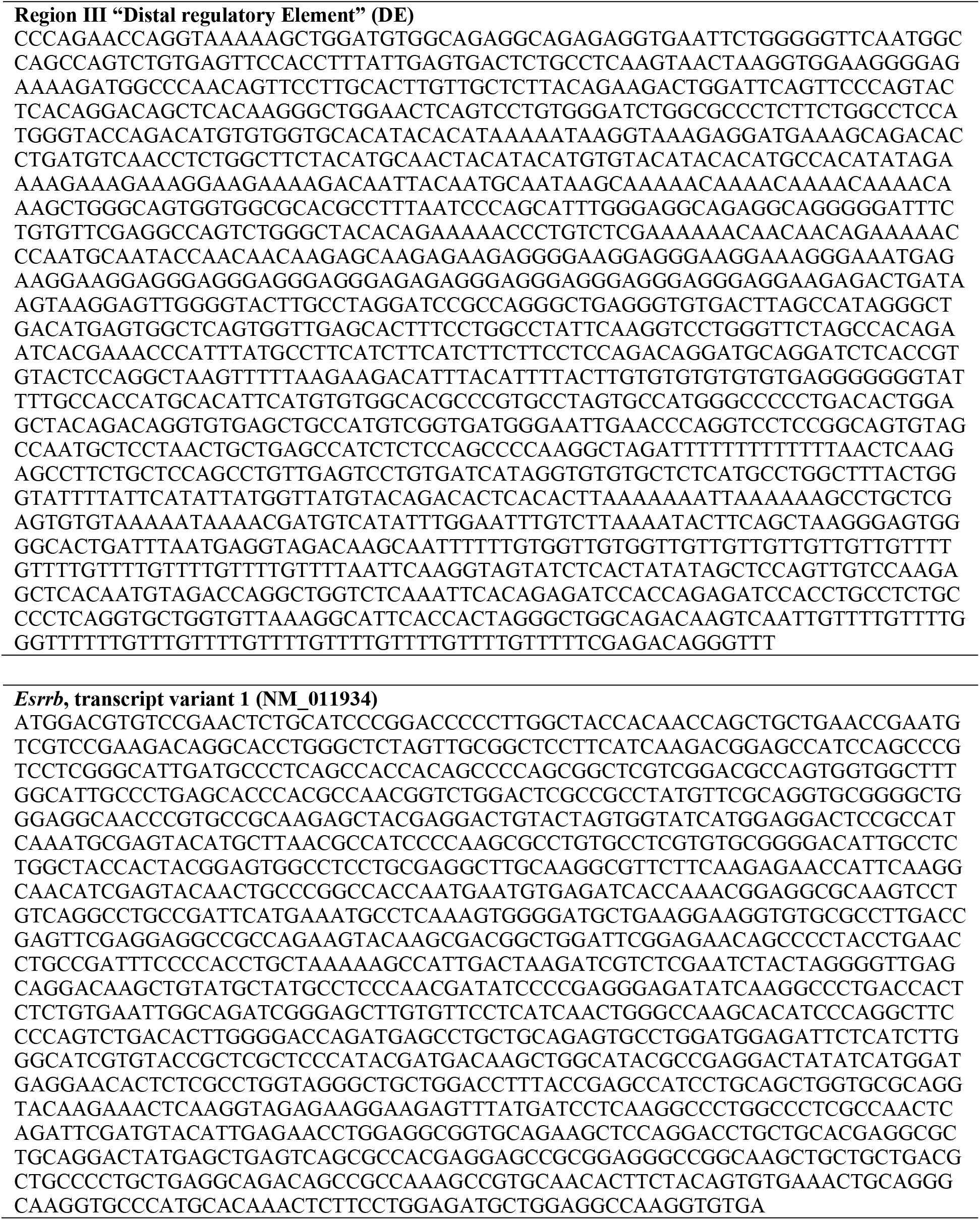
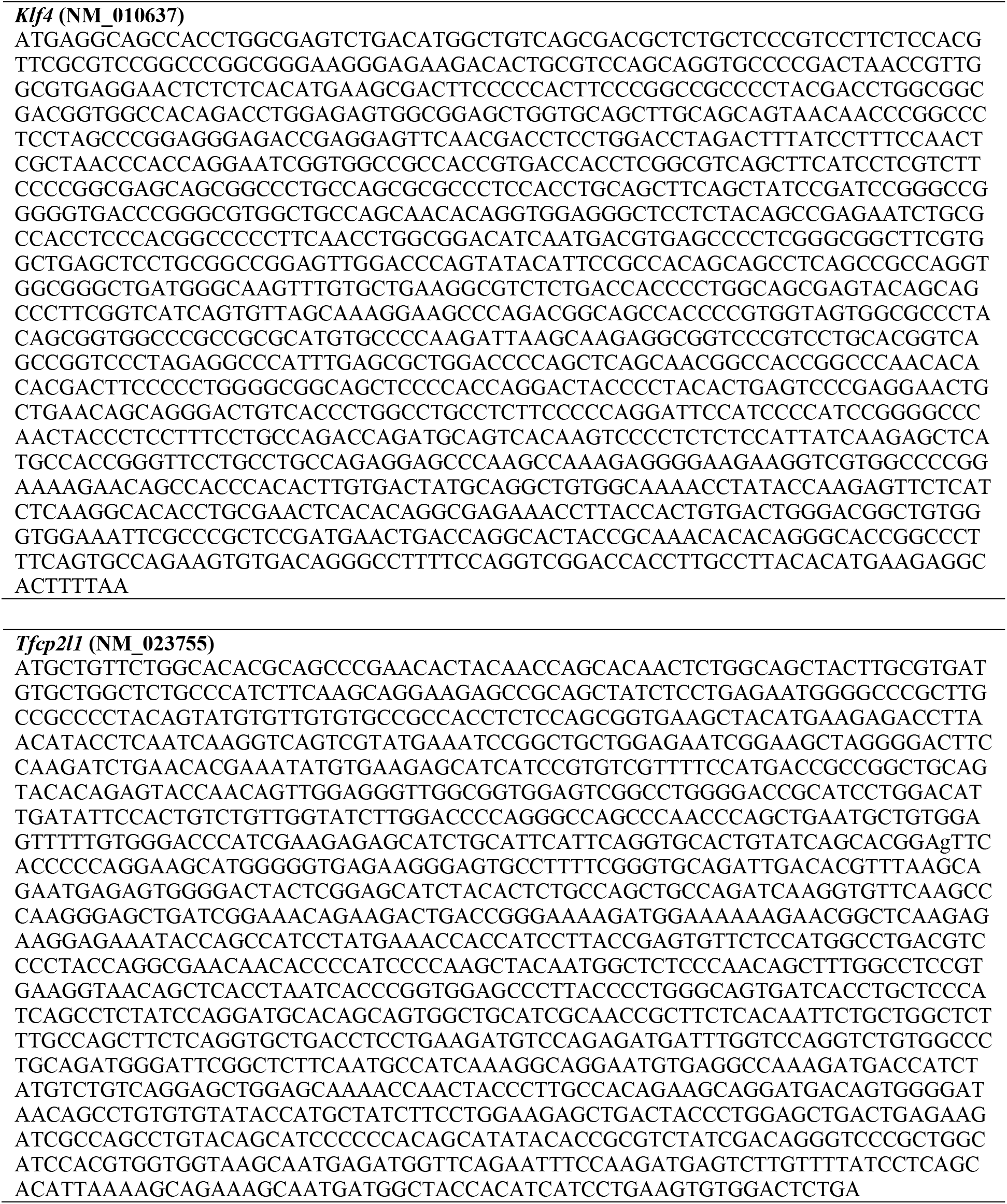
Nucleotide sequences (from 5’ to 3’) of the *Cyclin E* promoter regions I (P), II (PE) and III (DE), synthetized by GeneArt (Invitrogen), and of the *Esrrb*, *Klf4* and *Tfcp2l1* CDS, cloned for the overexpression.

**Table S4:**
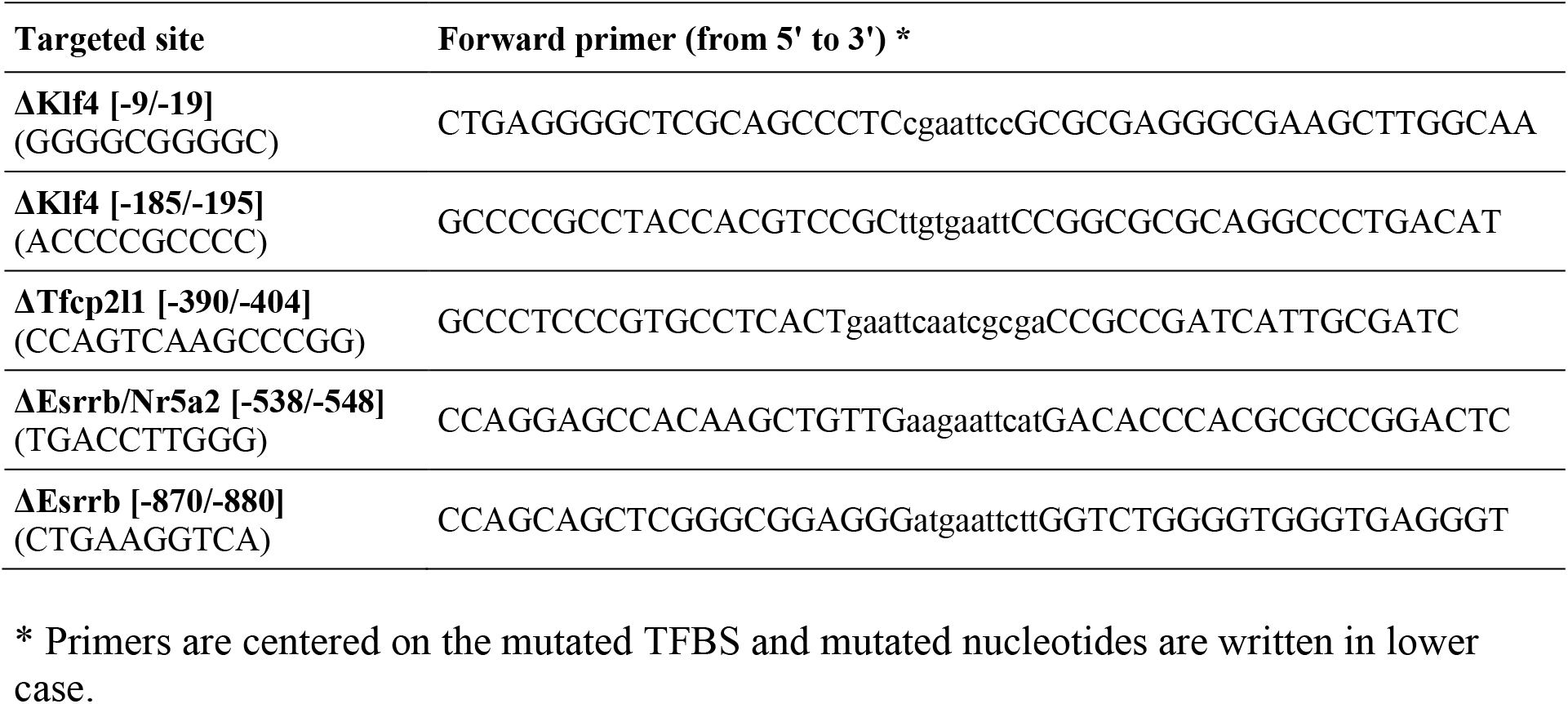
Primers designed for site-directed mutagenesis of ESRRB, KLF4 and TFCP2L1 binding sites.

**Table S5:**
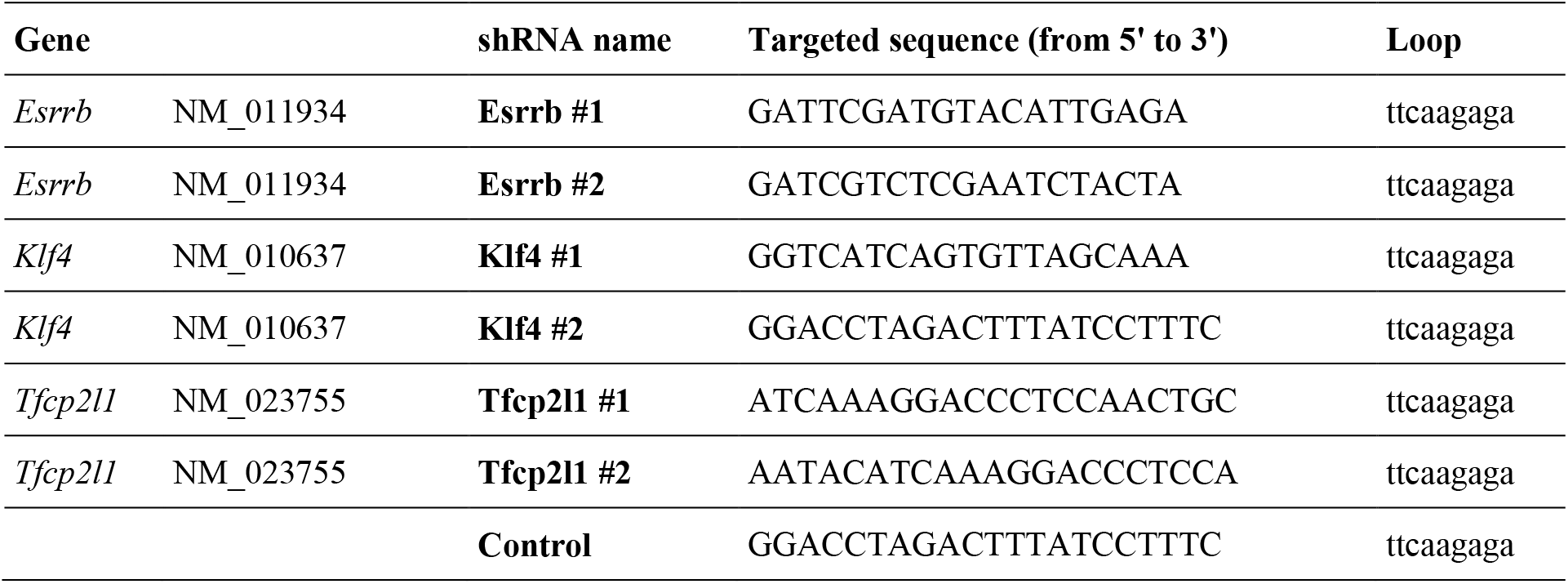
shRNA sequences.

